# A rare pre-existing progenitor-like Primed SMC compartment is the dominant inferred source of SMC-derived cellularity in vascular injury and atherosclerosis

**DOI:** 10.64898/2026.06.28.735042

**Authors:** Shwethali Wani, Michael Kitching, Sobhan Aboulhassanzadeh, Teodora-Simona Lungu, Isil Kilicgun, Karsyn Ulibarri, Weimin Liu, Achilleas Floudas, Eileen M Redmond, Paul A Cahill

## Abstract

The contribution of pre-existing smooth muscle cell (SMC) states to vascular lesions remains unresolved. Here we identify a rare, progenitor-like Primed SMC compartment before lesion formation in sham-operated carotid arteries and across *Myh11*-lineage–marked mouse aortic datasets, with a Primed-like counterpart in non-diseased human aorta. Primed SMCs retain SMC identity while combining attenuated contractile programmes with conserved niche-associated (*Cd34, Fst, Tnfrsf11b*) and matricellular (*Vcam1, Thbs1, Timp1*) cores, overlaid by vessel-specific developmental signatures (*Nrn1, Gdf10, Scx*). After carotid ligation and during high-fat diet–induced atherosclerosis in *Ldlr*^−/−^ and *Apoe*^−/−^ mice, disease-associated Primed, cycling and fibrochondrocyte populations preferentially map to baseline Primed SMCs, whereas contractile SMCs are depleted as inferred feeder sources. Concordant trajectory, graph-based and cross-dataset analyses support a model in which this pre-existing compartment is the dominant inferred source of SMC-derived lesional cellularity. These findings favour expansion and diversification of the Primed compartment over widespread phenotypic switching of contractile SMCs.

## Main text

Atherosclerosis is a leading cause of myocardial infarction and stroke, and smooth muscle cell (SMC)-derived cells constitute 40–70% of advanced lesion cellularity^1^. The prevailing model holds that medial contractile SMCs switch phenotype, lose contractile markers and acquire fibroblast- or macrophage-like identities^2–4^. Myh11-CreERT² tracing showed extensive SMC modulation, and SMC-specific *Klf4* deletion reduced lesion size and increased cap thickness^5,6^. Multicolour tracing further showed oligoclonal accumulation from a restricted subset of Myh11-expressing medial cells whose progeny span lesion phenotypes^7–10^. The constitutive identity of this subset and its relationship to vessel-wall progenitors remain unresolved.

Healthy-vessel single-cell analyses identified rare Sca1⁺/*Ly6a*⁺ SMCs (0.2–1.6% of medial cells) expressing disease-associated programmes^10,11^. Yet Sca1-CreERT² mainly marks adventitial rather than medial SMCs and contributes little to lesions^12,13^. *Gli1*⁺ adventitial cells^14–17^ and *Sox10*⁺/*S100b*⁺ vascular progenitors^18–21^ also show context-dependent contributions, but their relationship to constitutive Myh11-lineage subpopulations is unclear. Thus, atlas studies repeatedly identify Sca1⁺ disease populations that pre-injury Sca1 tracing scarcely labels. This discrepancy has not been tested directionally against the cell states available before and during disease in independent models.

Here we analyse carotid ligation, healthy Myh11-lineage-traced aortic media, longitudinal *Ldlr*^−/−^ and *Apoe*^−/−^ atherosclerosis, and healthy human aorta. A rare Primed SMC compartment is present at homeostasis in every model, retains SMC identity and combines attenuated contractile programmes with matricellular, niche/progenitor and chondrogenic-poised programmes. kNN feeder-source mapping, tested against the reference pools available at each stage and supported by pseudotime, rooted-tree, graph and temporal-flow analyses, identifies a common architecture: autonomous Primed expansion, depletion of contractile SMCs as a feeder source, and a Primed origin for cycling and fibrochondrocyte populations. Baseline recovery across cohorts tests continuity with a constitutive state rather than retrospective similarity to a disease-emergent population. The data favour expansion of a pre-existing compartment over widespread switching of contractile SMCs and nominate intersectional Myh11-lineage tracing gated on *Cd34* or *Vcam1* as a decisive test.

## Results

### A rare Primed SMC compartment is present in the healthy carotid artery

Single-cell RNA sequencing (scRNA-seq) analysis of murine carotid arteries 14 days after sham operation or ligation used 10 carotid arteries from 10 mice pooled before dissociation into one composite 10x library per condition. After QC, 1,973 sham and 4,272 ligated cells were retained, resolving 17 Harmony-defined clusters (Fig. 1A,B; Supplementary Table 1). Sham arteries were dominated by quiescent contractile SMCs (1,534/1,973 cells; 77.7%) but also contained a smaller, transcriptionally distinct SMC population present at homeostasis, hereafter termed Primed SMCs (101/1,973; 5.1%).

**Fig. 1.**
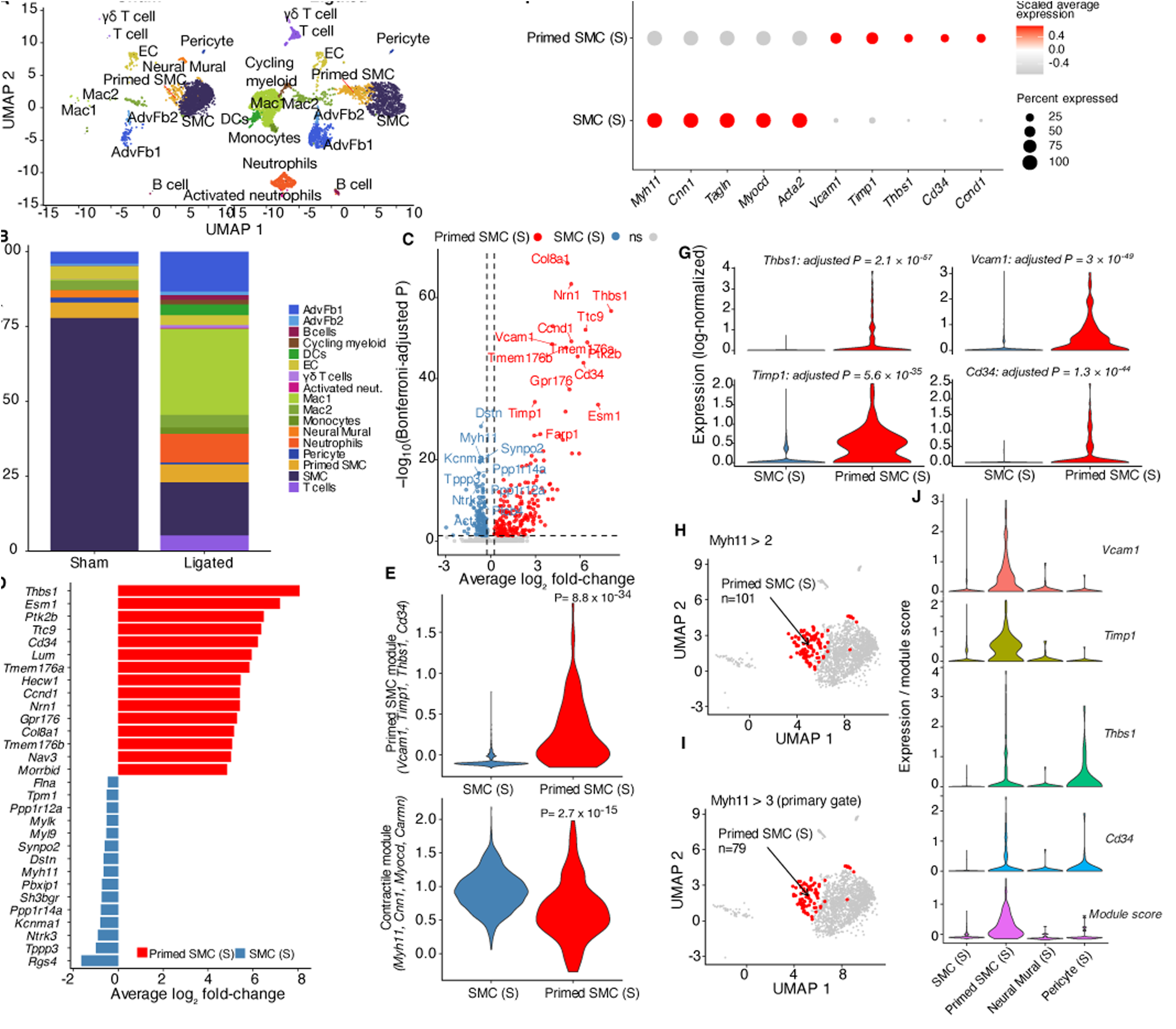
A rare Primed SMC compartment is present in the healthy carotid artery and shows a modest increase in relative representation after ligation. **(A)** Harmony-integrated UMAP of scRNA-seq from murine carotid arteries 14 d after sham operation or ligation. Ten vessels from 10 mice were pooled into one library per condition (sequencing n = 1 pool per condition); 1,973 sham and 4,272 ligated cells passed QC. Seventeen clusters were annotated using canonical markers (Supplementary Table 1). **(B)** Descriptive cell-type composition in the single pooled sham and ligated libraries. The contractile-SMC fraction decreased, whereas the fractions of Mac1/Mac2 macrophages, AdvFb1 fibroblasts and Primed SMCs increased; additional disease-associated populations were detected after ligation. These recovered-cell proportions were not tested across biological replicates. **(C)** Volcano plot, sham Primed SMC versus sham contractile SMC (n = 101 and 1,534 cells; two-sided cell-level Wilcoxon rank-sum test; Seurat Bonferroni correction across 24,527 assayed features). FindMarkers was run with logfc.threshold = 0; significance was defined post hoc as Bonferroni-adjusted P < 0.05 and |average log₂FC| > 0.25. Red, enriched in Primed SMCs; blue, enriched in contractile SMCs; grey, not significant; selected genes labelled. **(D)** Descriptive display of 15 Primed-enriched genes selected by decreasing average log₂FC and 15 contractile-SMC-enriched genes selected by smallest Bonferroni-adjusted P; bars show average log₂FC. **(E)** Primed-module score (*Vcam1, Timp1, Thbs1, Cd34*; upper, unadjusted P = 8.8 × 10⁻³⁴) and contractile-module score (*Myh11, Cnn1, Myocd, Carmn*; lower, unadjusted P = 2.7 × 10⁻¹⁵) in sham contractile versus Primed SMCs (n = 1,534 and 101 cells; separate two-sided cell-level Wilcoxon rank-sum tests). Scores are background-corrected relative values calculated with AddModuleScore; negative values do not denote absent expression. **(F)** Dot plot of representative contractile and Primed-associated genes in sham contractile versus Primed SMCs (dot size, percentage expressing; colour, scaled average expression). **(G)** Per-gene violins for the four Primed-module genes: *Thbs1*, Bonferroni-adjusted P = 2.1 × 10⁻⁵⁷; *Vcam1*, 3.0 × 10⁻⁴⁹; *Timp1*, 5.6 × 10⁻³⁵; and *Cd34*, 1.3 × 10⁻⁴⁴. Values are from the same cell-level differential-expression analysis as C. **(H)** UMAP of the 101 sham Primed SMCs. **(I)** The 79 sham Primed SMCs passing the *Myh11* > 3 log-normalised gate (red; 78.2%) on the full sham UMAP (grey); the gate is a transcriptional SMC-identity proxy. **(J)** Primed-module score violins in the *Myh11*-gated population, showing Primed-SMC-specific enrichment relative to contractile SMCs, neural mural cells and pericytes. The cell-level tests in C, E and G describe state separation within one pooled sham library and do not constitute biological-replicate inference.

Primed SMCs displayed progenitor-niche features but lacked a classical stem-cell identity (Fig. 1C,D; Extended Data Fig. 1). They expressed matricellular and ECM genes (*Thbs1, Lum, Dcn, Col8a1*), niche-associated genes (*Cd34, Cxcl12, Tnfrsf11b, Fst*), developmental genes (*Nrn1, Gdf10, Scx*) and *Ccnd1*. This programme enriched vascular development, endothelial-cell migration, SMC proliferation and biomineral-tissue development terms and PI3K–Akt, ECM–receptor, AGE–RAGE and Rap1 signalling pathways, but not canonical stem-cell markers: *Ly6a*/Sca1 and *Kit* were absent or low, and *Sox2, Nanog* and *Pou5f1* were undetectable. Within the pooled sham library, Primed SMCs had higher Primed-module scores and lower contractile-module scores than contractile SMCs (unadjusted two-sided cell-level Wilcoxon P = 8.8 × 10⁻³⁴ and 2.7 × 10⁻¹⁵, respectively), while retaining attenuated contractile-gene expression (Fig. 1E-G; Extended Data Fig. 2a). These tests quantify transcriptional separation among captured cells and do not estimate between-animal variability. Seventy-nine of 101 cells (78.2%) retained *Myh11* >3 while co-expressing the Primed signature; other Myh11-high mural populations did not (Fig. 1H-J; Extended Data Fig. 2b).

Independent de novo re-clustering of sham cells recovered the same compartment as a high-purity orthocluster (n = 56; 92.9% Primed; r = 0.97 with integrated Primed SMCs; Extended Data Fig. 3a–g) retaining contractile identity and the full Primed signature (Extended Data Fig. 3h), confirming that it is not an integration artefact. Relative to candidate vessel-wall progenitors, *Ly6a*/Sca1 and *S100b* were primarily concentrated in endothelium and adventitial fibroblasts. Primed SMCs contained only a minor *S100b*⁺ subset (∼7%) and were essentially *Ly6a*-negative (Extended Data Fig. 4a–g).

### Primed SMC relative representation increases modestly and a cycling Primed-like subset is detected after ligation

After ligation, contractile SMCs accounted for 17.8% of recovered cells versus 77.7% in sham, whereas Primed SMCs accounted for 6.0% (255/4,272) versus 5.1% (101/1,973) (Fig. 1A,B; Extended Data Fig. 2c,d). Because each condition was represented by one pooled library and total cell recovery differed, these changes are descriptive and do not establish absolute population expansion or a between-animal effect. Recovered contractile SMCs showed lower UMI counts and gene detection (−27% and −14%), whereas these cell-level QC metrics did not differ materially for Primed SMCs (UMI +8%, unadjusted two-sided Wilcoxon P = 0.18; genes +4%, P = 0.22; Supplementary Fig. 1a–c). These QC comparisons do not suggest a large loss of transcript complexity among captured Primed SMCs, but they cannot exclude condition-dependent differences in dissociation or cell recovery. The lower recovered contractile fraction is consistent with documented medial SMC apoptosis after ligation^22–24^, although the pooled design cannot distinguish biological loss from differential recovery.

Sub-clustering of the residual contractile compartment identified five sub-states, dominated by an injury-emergent stress state. However, Primed module scores remained near baseline in every contractile sub-cluster (Extended Data Fig. 5a-e). In sham samples, *Cd34* was strongly enriched in Primed SMCs relative to contractile SMCs (+6.2 log₂FC; 25.7% versus 1.2%). Among captured Primed SMCs, *Cd34* expression was similar in the sham and ligated libraries and remained low across all contractile SMC subclusters (0-7%). This descriptive pattern is consistent with preservation of a *Cd34*⁺ Primed state after ligation; it does not by itself demonstrate expansion or exclude recruitment from another state.

Within the single pooled ligated library, 20 Primed SMCs met the operational Cycling Primed SMC definition: assignment to S or G2/M phase together with detectable expression (counts > 0) of each of Mki67, Top2a and Birc5. These cells occupied a region adjacent to Primed SMCs and distinct from cycling myeloid cells (Fig. 2A; Extended Data Fig. 6a-c). As expected from their definition, they had higher mitotic-marker expression and cycling-module scores than contractile and non-cycling Primed SMCs (Extended Data Fig. 6d-f). They retained *Vcam1, Timp1* and *Thbs1*, and their Primed and contractile module scores did not differ detectably from those of non-cycling Primed SMCs (Fig. 2B-D). These findings identify a cycling Primed-like transcriptional state but do not establish its lineage origin or exclude convergence from contractile SMCs.

**Fig. 2.**
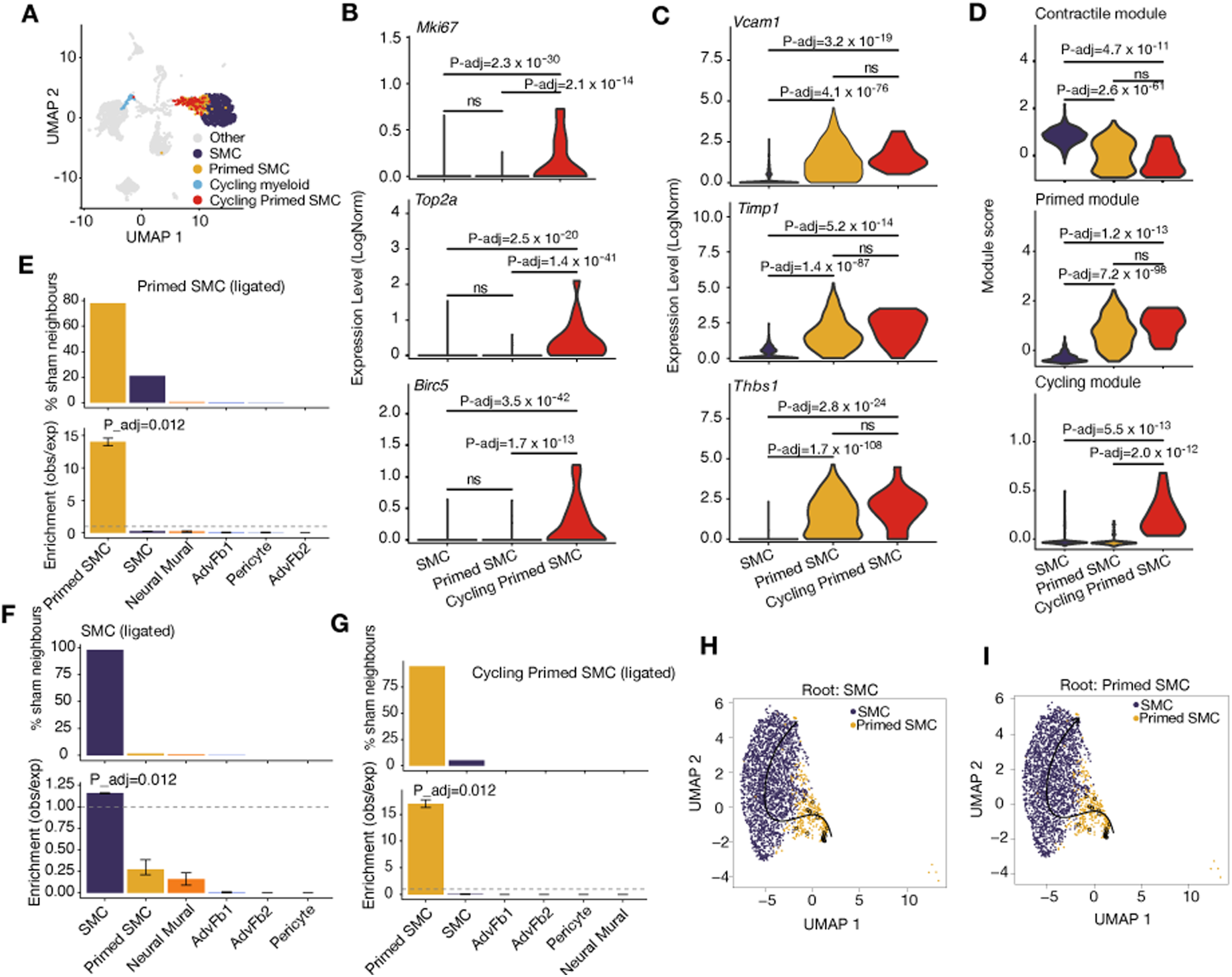
Cycling Primed SMCs retain the Primed programme and map preferentially to sham Primed SMCs after carotid ligation. **(A)** UMAP of 4,272 cells from the single pooled ligated carotid library. Contractile SMCs (labelled SMC; n = 761), non-cycling Primed SMCs (n = 235), Cycling Primed SMCs (n = 20) and Cycling myeloid cells (n = 70) are highlighted; the remaining cells are shown in grey. Cycling Primed SMCs were defined as ligated Primed SMCs assigned to S or G2/M phase and with detectable expression (counts > 0) of each of Mki67, Top2a and Birc5. **(B)** Per-cell log-normalized expression of *Mki67, Top2a* and *Birc5* in ligated contractile SMCs, non-cycling Primed SMCs and Cycling Primed SMCs. **(C)** Per-cell log-normalized expression of *Vcam1, Timp1* and *Thbs1* in the same groups. **(D)** Contractile (*Myh11, Cnn1, Myocd* and *Carmn*), Primed (*Vcam1, Timp1, Thbs1* and *Cd34*) and cycling (*Mki67, Top2a* and *Birc5*) module scores calculated using AddModuleScore. Module scores are background-corrected relative values; negative values do not denote absent expression. For B-D, groups comprised 761 contractile SMCs, 235 non-cycling Primed SMCs and 20 Cycling Primed SMCs. Comparisons used separate two-sided cell-level Wilcoxon rank-sum tests, with Bonferroni correction across the three pairwise group comparisons within each gene or module; ns, adjusted P ≥ 0.05. **(E-G)** Forward k-nearest-neighbour mapping of ligated target cells to a sham mesenchymal reference comprising 1,812 contractile SMCs, Primed SMCs, AdvFb1 cells, AdvFb2 cells, neural mural cells and pericytes. For each target cell, the k = 15 nearest sham neighbours were identified in 20-dimensional Harmony space. Upper bars show the percentage of all neighbour assignments contributed by each sham reference population. Lower bars show observed/expected enrichment relative to that population’s frequency in the sham reference; the dashed line denotes an enrichment of 1. Error bars show 95% percentile confidence intervals from 50 bootstrap resamples of target cells. The plotted adjusted P value refers to the most enriched reference population and was calculated using a one-sided, Laplace-corrected test based on 500 reference-label permutations, with Bonferroni correction across the six reference populations. **(E)** All ligated Primed SMCs (n = 255, including the 20 cycling cells) mapped predominantly to sham Primed SMCs (77.9% of neighbour assignments; 14.0-fold enrichment; adjusted P = 0.012); sham contractile SMCs contributed 21.2% of neighbours and were under-represented relative to their reference frequency (0.25-fold). **(F)** Ligated contractile SMCs (n = 761) mapped predominantly to sham contractile SMCs (98.0%; 1.16-fold enrichment; adjusted P = 0.012); sham Primed SMCs contributed 1.5% of neighbours (0.27-fold). **(G)** Cycling Primed SMCs (n = 20) mapped predominantly to sham Primed SMCs (94.7%; 17.0-fold enrichment; adjusted P = 0.012); sham contractile SMCs contributed 5.3% of neighbours (0.063-fold). **(H,I)** Dual-rooted Slingshot analysis of the combined sham and ligated SMC axis, comprising 2,295 contractile and 356 Primed SMCs. Curves were fitted in 20-dimensional Harmony space and projected onto UMAP, with the root placed at contractile SMCs in H or Primed SMCs in I. Reversing the root reverses the inferred pseudotime direction, illustrating that Slingshot does not independently determine biological direction. The carotid experiment comprised one pooled library per condition, each generated from 10 carotid arteries from 10 mice. All n values denote captured cells. Cell-level tests and nearest-neighbour mapping describe cells within or between the two pooled libraries and do not estimate between-animal variability or establish clonal ancestry.

Within the pooled libraries, ligated Primed SMCs showed higher *Ly6a* expression than sham Primed SMCs (Bonferroni-adjusted cell-level P = 7.4 × 10⁻⁸), no detectable change in *S100b* and lower *Myh11* expression (BH-adjusted cell-level P = 2.0 × 10⁻³²; Extended Data Fig. 4h-j). These comparisons describe captured cells within the two pooled libraries.

### Ligated Primed and cycling states preferentially map to sham Primed SMCs

To assess which pre-injury state each ligated population most closely resembled, we mapped each ligated cell to its 15 nearest sham neighbours in 20-dimensional Harmony space and compared the observed neighbour composition with the abundance of each sham reference population. For all 255 ligated Primed SMCs, 77.9% of neighbour assignments were sham Primed SMCs, corresponding to 14.0-fold enrichment over expectation (Bonferroni-adjusted permutation P = 0.012); sham contractile SMCs were under-represented (0.25-fold; Fig. 2E). Ligated contractile SMCs mapped predominantly to sham contractile SMCs (98.0% of neighbours), whereas sham Primed SMCs were under-represented (0.27-fold; Fig. 2F). The 20 Cycling Primed SMCs also mapped predominantly to sham Primed SMCs (94.7%; 17.0-fold enrichment; adjusted P = 0.012; Fig. 2G). Ligated AdvFb1 similarly mapped to sham AdvFb1, whereas ligated Mac1 mapped to sham myeloid populations (Extended Data Fig. 6g,h), supporting the cell-state specificity of the procedure. Bootstrap intervals reflect resampling of captured cells, and permutation P values reflect reference-cluster label shuffling; neither estimates between-mouse variability. These mappings support transcriptional continuity with the sham Primed state but do not demonstrate clonal ancestry or absolute population expansion.

Dual-rooted Slingshot placed contractile and Primed SMCs on the same connected trajectory, but reversing the selected root reversed the inferred pseudotime direction (Fig. 2H,I). The topology is therefore compatible with transcriptional continuity but does not determine which state precedes the other. In the reciprocal analysis, which queried sham cells against ligated neighbours, ligated Primed SMCs were under-represented among the neighbours of sham contractile SMCs (0.22-fold; Extended Data Fig. 7a–c). This reciprocal pattern is concordant with the forward mapping but is not an independent lineage test. Both nearest-neighbour mapping and Slingshot quantify transcriptomic relationships rather than clonal ancestry and cannot exclude convergence of modulated contractile SMCs on a Primed-like state. We therefore examined two genetic lineage-tracing models of atherosclerosis, in which temporal kinetics could further test the inferred state relationships.

### Autonomous expansion of the Primed SMC pool dominates lesion cellularity in atherosclerosis

We analysed Myh11-CreERT² ZsGreen1-WPRE lineage-traced *Ldlr*^−/−^ and *Apoe*^−/−^ datasets spanning Control, Weeks 8 and 16, and late disease (Week 26 *Ldlr*^−/−^; Week 22 *Apoe*^−/−^)^25^. A ZsGreen1-WPRE >3.17 gate, restricted to SMC-lineage compartments, retained 11,362 *Ldlr*^−/−^ and 10,966 *Apoe*^−/−^ lineage-positive cells. Harmony clustering resolved contractile and Primed SMCs, adventitial fibroblasts, pericytes, a fibromyocyte/fibrochondrocyte (FbC) compartment, cycling cells and a minor macrophage-like population (Fig. 3A; Supplementary Table 2).

**Fig. 3.**
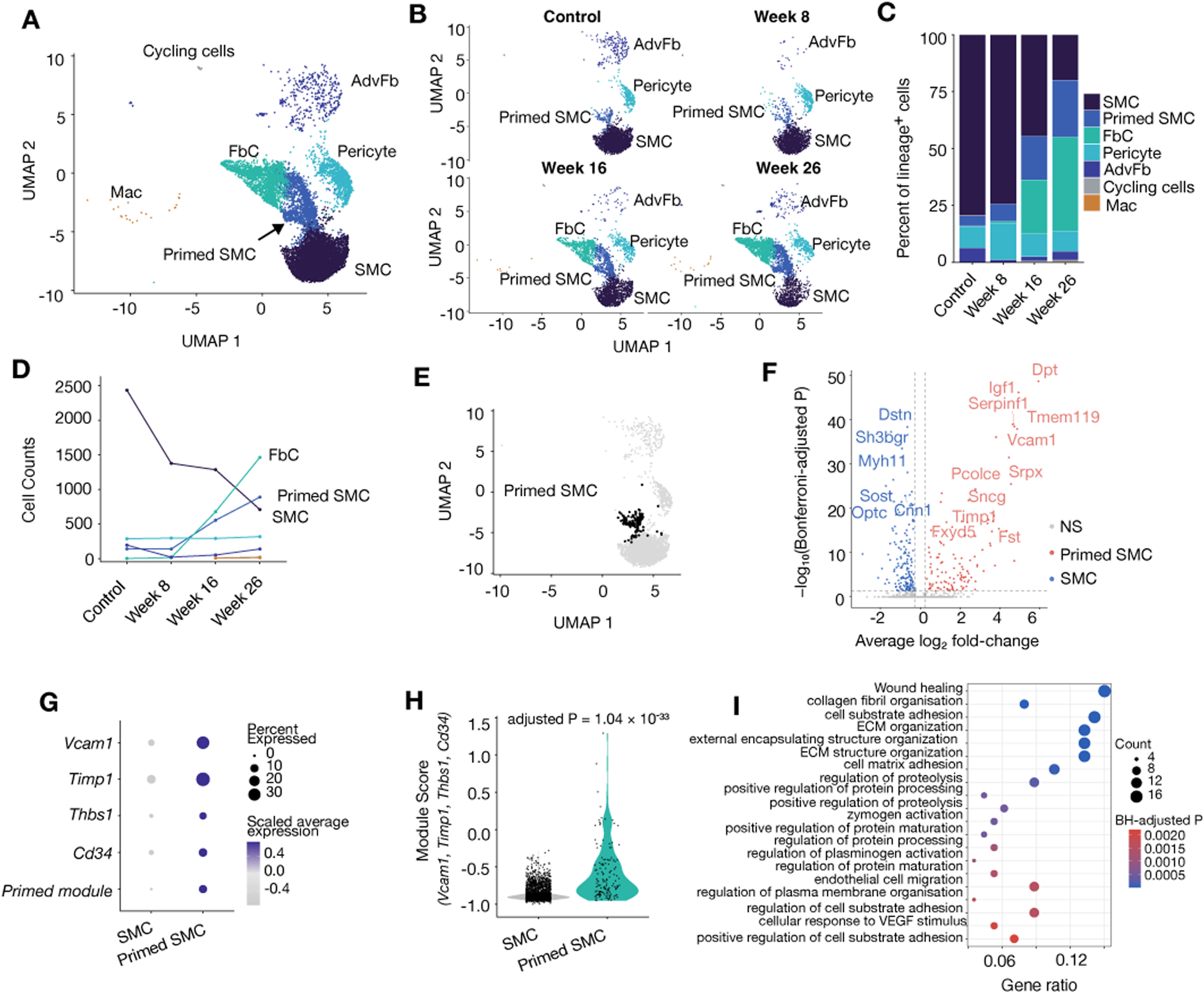
A pre-existing Primed SMC pool expands progressively in *Ldlr*^−/−^ atherosclerosis. *Ldlr*^−/−^ Myh11-CreERT² ZsGreen1-WPRE scRNA-seq; lineage-positive cells retained at ZsGreen1-WPRE >3.17 log-normalised counts (n = 11,362). (A) Harmony-integrated UMAP showing contractile SMC, Primed SMC, FbC, AdvFb, Pericyte/Neural Mural, Cycling cells and Mac. (B) UMAP by time point. (C) Lineage-positive composition across Control and Weeks 8/16/26. (D) Cell counts: contractile SMC decline 3.4-fold, Primed SMC expand 6.3-fold and FbC become the largest compartment by Week 26. (E) Control Primed SMC on the UMAP. (F) Volcano plot, Control Primed versus contractile SMC. (G) Primed-module dot plot. (H) Primed-module score in Control SMC versus Primed SMC (p_adj = 1.04 × 10⁻³³). (I) GO Biological Process enrichment among Control Primed-enriched genes.

The *Ldlr*^−/−^ lineage-positive compartment underwent marked remodelling (Fig. 3B–D; Supplementary Fig. 2a–d). Contractile SMCs declined from 79.5% at Control to 19.9% at Week 26 (2,435 to 710 cells; 3.4-fold), whereas Primed SMCs increased from 4.6% to 25.0% (141 to 890 cells; 6.3-fold). FbC were almost absent at Control and Week 8 (3 and 15 cells) but emerged by Week 16 (679 cells; 23.5%) and became the largest lineage-positive population at Week 26 (1,464 cells; 41.1%). Together, Primed SMCs, FbC and cycling cells increased from 4.7% to 66.8%. By Week 8, the SMC compartment had already contracted by 41%, driven by a 43% loss of contractile SMCs, whereas Primed numbers were unchanged (141 versus 140) and FbC remained rare. Thus, contractile loss preceded Primed and FbC expansion. The lineage-positive fraction remained stable (Supplementary Fig. 2e,f), arguing against reporter silencing. Early contractile decline is consistent with documented medial SMC apoptosis^24,26,27^. Subsequent recovery of the broader lineage compartment reflected Primed expansion and FbC emergence rather than contractile restoration. Control Primed SMCs preserved the homeostatic identity (Fig. 3E–H) and enriched ECM and collagen organisation, adhesion, wound healing, proteolysis and plasminogen activation (Fig. 3I).

### Temporal feeder analysis supports autonomous expansion of Primed SMC during atherosclerosis

Temporal kNN analysis queried each time point against all preceding cells, with bootstrap resampling and permutation testing (Fig. 4A). Primed self-feeding increased from 73.1% at Week 8 to 76.7% at Week 16 and 88.0% at Week 26 (p_adj ≤ 0.014; Fig. 4B). The declining enrichment values (15.9-to 8.2-fold) reflected growth of the Primed fraction in the reference rather than loss of specificity. Contractile SMCs remained below background as a Primed source (0.30-, 0.21- and 0.07-fold). At Week 16, the largest baseline-derived FbC contribution came from Primed SMCs (35.9%; 6.3-fold), with negligible contractile input; cycling cells also mapped mainly to Primed SMCs (58.8%) with no detected contractile input. By Week 26, FbC were predominantly self-maintaining, whereas the small cycling population showed k-sensitive self-renewal (Fig. 4C–F; Supplementary Fig. 3c).

**Fig. 4.**
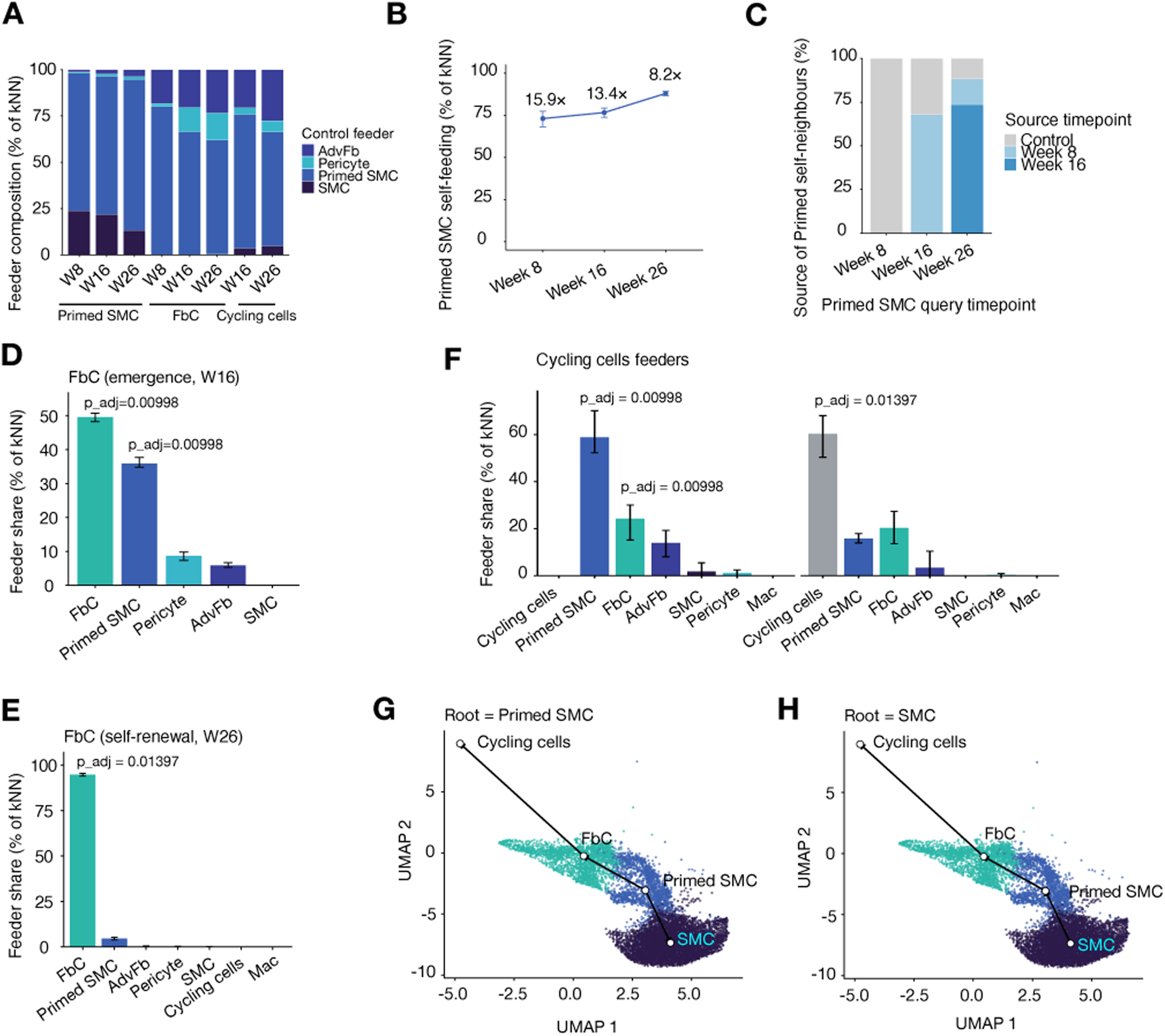
Temporal feeder-source analysis identifies the Primed pool as the dominant inferred source in *Ldlr*^−/−^ atherosclerosis. Forward kNN analysis of lineage-positive cells across Control and Weeks 8/16/26 (20 Harmony dimensions, k = 15); error bars, bootstrap 95% CI; P, 500-permutation test with Bonferroni correction. (A) Baseline neighbour composition of each query. (B) Primed self-feeding rises from 73.1% to 88.0%, while contractile SMCs are progressively depleted as feeders. (C) Source-time-point decomposition shows stepwise propagation through preceding Primed pools. (D) FbC at emergence map chiefly to existing FbC and Primed SMCs, with negligible contractile input. (E) Week-26 FbC are predominantly self-maintaining. (F) Cycling cells map mainly to Primed SMCs at emergence; the small Week-26 population shows k-sensitive self-renewal. (G,H) Dual-rooted Slingshot over the SMC-fate compartment, rooted at Primed (G) or contractile SMC (H). Slingshot establishes topology; direction is adjudicated by the feeder analysis.

Rooted-tree analysis placed contractile SMCs at a terminal node and Primed SMCs at an internal node. With a Primed root, the analysis resolved two lineages (Primed → FbC → Cycling and a branch to SMC; Fig. 4G). A contractile root instead forced a single trajectory (SMC → Primed → FbC → Cycling; Fig. 4H). The topology therefore does not require a contractile root and is compatible with, but does not itself prove, Primed SMCs occupying the apex.

The independently analysed *Apoe*^−/−^ model reproduced baseline Primed module separation (p_adj = 1.37 × 10⁻²⁴; Fig. 5G), Primed expansion and the emergence and dominance of FbC retaining the Primed programme (Fig. 5A–I; Supplementary Table 3). Shared features included ECM, adhesion, vascular-remodelling and ossification enrichment; Primed self-feeding, which declined from 23.6-fold at Week 8 to 8.5-fold at Week 22; contractile depletion; contributions from Primed SMC and AdvFb to FbC; and equivalent dual-rooting (Extended Data Fig. 8a–h). Cell-cycle regression removed an apparent contractile contribution to cycling cells (Supplementary Fig. 3a,b,d). Injury-associated contractile subclusters in both models retained contractile programmes and did not acquire the defining Primed markers (Extended Data Fig. 5f–o). Alongside an independent healthy aortic-medial dataset, these formed the four baseline cohorts used below.

**Fig. 5.**
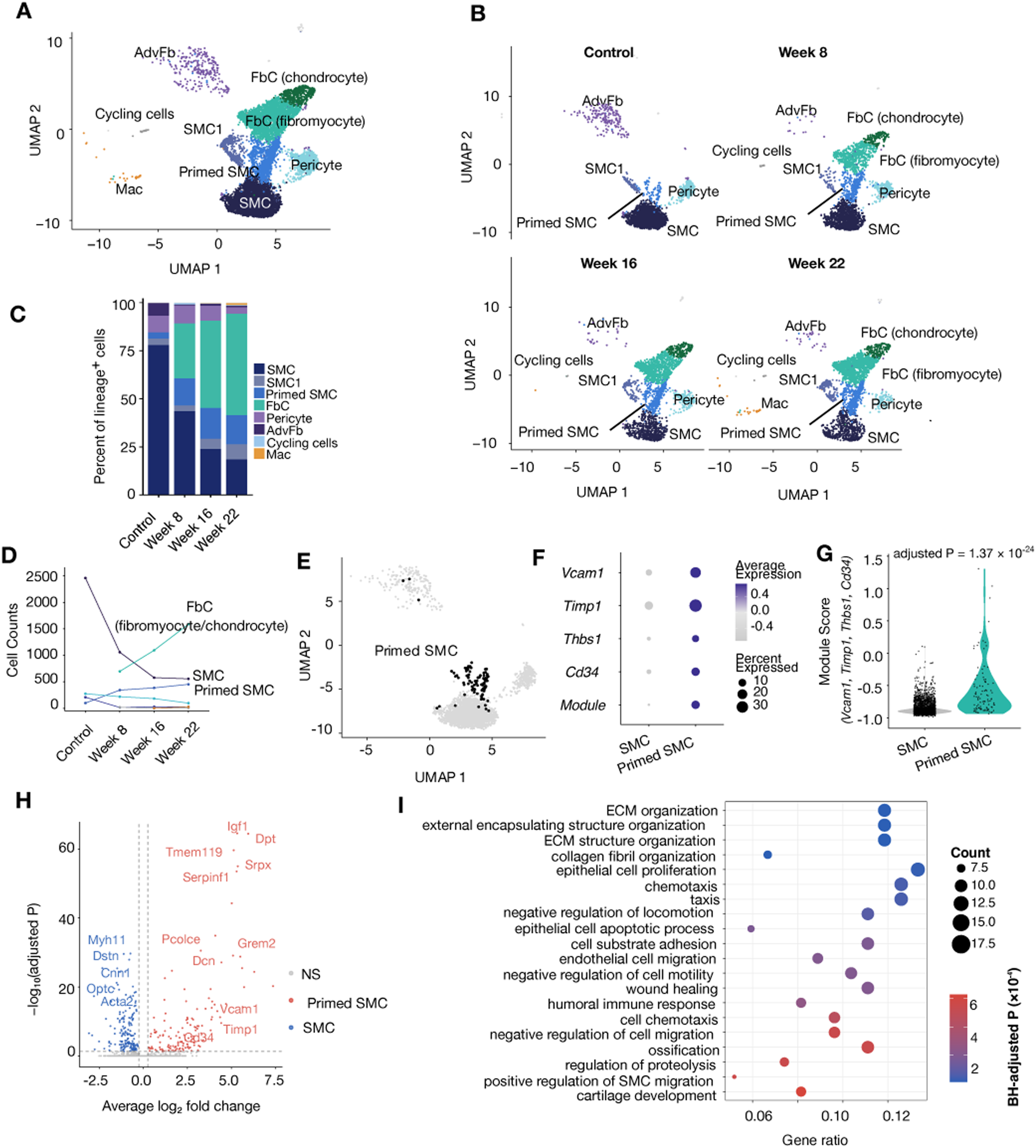
*Apoe*^−/−^ atherosclerosis independently recapitulates the autonomous-expansion architecture. Myh11-CreERT²;ZsGreen1-WPRE lineage-traced *Apoe*^−/−^ aortic cells across Control and Weeks 8/16/22; lineage-positive cells (ZsGreen1-WPRE >3.17; n = 10,966) clustered independently of *Ldlr*^−/−^. (A) UMAP by cell type. (B) UMAP by time point. (C) Lineage-positive composition. (D) Counts showing reciprocal contractile loss and Primed/FbC expansion. (E) Control Primed SMC on the UMAP. (F) Primed-marker and module dot plot. (G) Primed-module score, contractile versus Primed SMC (p_adj = 1.37 × 10⁻²⁴). (H) Volcano plot, Control Primed versus contractile SMC. (I) GO Biological Process enrichment among Primed-enriched genes.

### *Ly6a* and *S100b* mark the pre-existing Primed SMC pool and its disease-induced activation

Cross-cohort comparison at baseline using BH-corrected Wilcoxon tests showed that *S100b* and *Ly6a* mark complementary components of the Primed programme (Extended Data Fig. 4k,l). *S100b* was enriched in Primed SMCs across all four cohorts, reaching significance in sham carotid and aortic media, with borderline enrichment in *Ldlr*^−/−^ and a concordant trend in *Apoe*^−/−^. By contrast, *Ly6a* enrichment was largely restricted to aortic media, recapitulating the canonical Ly6a/Sca1⁺ aortic Primed state^10^ and showing that Ly6a is not a universal homeostatic marker. During disease, *Ly6a* was strongly induced in Primed SMCs in both models, whereas *S100b* remained relatively stable (Extended Data Fig. 4m,o). FbC expressed both markers from emergence and maintained strong *Ly6a* expression (Extended Data Fig. 4n,p), supporting molecular continuity with Primed SMCs. Neither marker alone defines the state; they represent overlapping subprogrammes within a broader conserved identity.

### Primed SMC carry a conserved *Cd34*⁺ progenitor-like programme as a single poised state

To define the intrinsic identity of Primed SMCs, we compared baseline Primed and contractile SMCs across four cohorts. Every dataset revealed a conserved niche/progenitor module (*Cd34, Tnfrsf11b, Phlda1, Fst, Eng*) and a matricellular module (*Vcam1, Thbs1, Timp1, Mgp, Lum, Dcn*; Extended Data Fig. 9e). The programme was progenitor-associated rather than pluripotent: canonical stem-cell markers (*Sox2, Nanog, Pou5f1, Kit*) were not enriched, *Ly6a* enrichment was confined to aortic media, and phenotypic-switching factors (*Klf4, Sox9, Nes*) were not enriched. *Runx2* increased only in carotid and aortic medial datasets, consistent with bed-specific chondrogenic priming. The contractile programme nevertheless remained robustly expressed. Niche/progenitor and matricellular modules defined a coupled state rather than distinct subpopulations: their scores correlated positively across all four cohorts (Spearman ρ = +0.26 to +0.58) and both tended to oppose the contractile programme (Extended Data Fig. 9a–d). Hartigan’s dip test found no bimodality (all p > 0.23), and sub-clustering found no progenitor-pure or contractile-pure populations. Primed SMCs therefore form a continuous progenitor-like state on a retained contractile background.

### A pan-vascular Primed SMC identity with bed-predominant axes

Using a common pipeline, we compared baseline Primed and contractile SMCs across all four cohorts (Fig. 6A). The healthy-aortic component of this comparison provided an independent test of conservation. Analysis of four replicate Myh11-CreERT² lineage-traced aortic medial libraries^10^ resolved a discrete Primed population alongside contractile SMCs (Fig. 6B,C). It comprised 240 lineage-marked cells at a consistent frequency across replicates (1.91–2.08%; Fig. 6D). Contractile markers (*Myh11, Cnn1, Tagln, Acta2*) remained high but modestly reduced, while *Vcam1, Timp1, Thbs1* and *Cd34* were selectively enriched (Extended Data Fig. 10a–c). Genome-wide differential expression identified 296 genes separating matricellular/ECM from contractile and oxidative programmes (Extended Data Fig. 10d,e), demonstrating conservation in healthy aortic media and reproducibility across libraries.

**Fig. 6.**
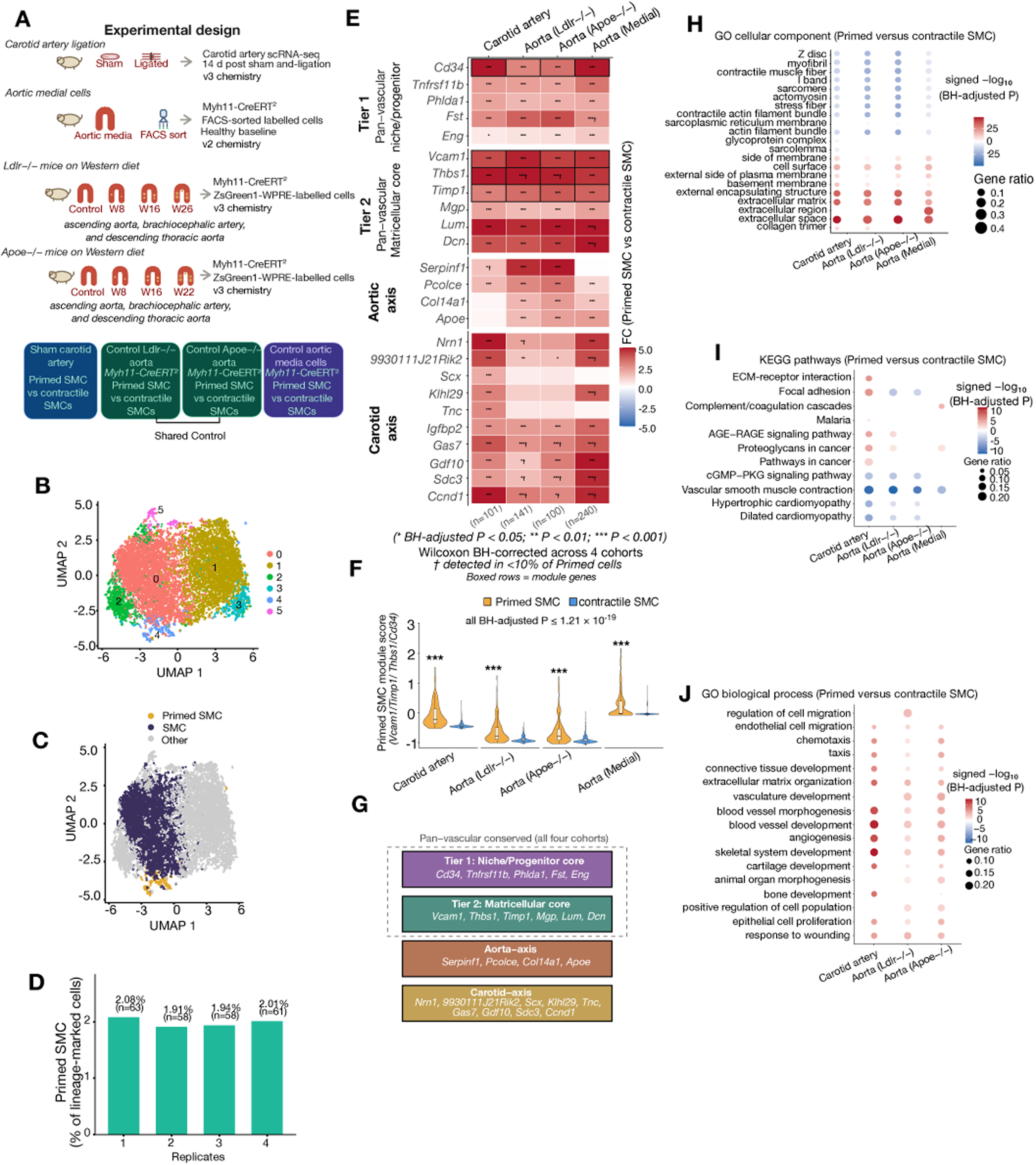
A conserved progenitor-like Primed SMC identity is shared across healthy vascular beds. (A) Design of the four baseline comparisons. (B–D) FACS-purified Myh11-CreERT² lineage-marked healthy aortic medial cells: UMAP by cluster (B), Primed and contractile SMC highlighted (C), and consistent Primed frequency across four replicate libraries (D; n = 63, 58, 58 and 61; 1.91–2.08%). (E) Tiered Primed-versus-contractile fold-change heatmap defining conserved niche/progenitor and matricellular cores and aorta- and carotid-predominant axes. (F) Four-gene Primed-module scores across cohorts. (G) Summary of conserved cores and bed-predominant axes. (H–J) GO Cellular Component (H), KEGG (I) and GO Biological Process (J) enrichment. Detailed healthy-aortic marker, differential-expression and enrichment analyses are shown in Extended Data Fig. 10.

Differential expression resolved two conserved cores—niche/progenitor and matricellular—and two bed-predominant axes: an aortic axis (*Serpinf1, Pcolce, Col14a1, Apoe*) and a carotid axis (*Nrn1, Scx, Tnc, Igfbp2, Gas7, Gdf10, Ccnd1*) (Fig. 6E). The prespecified four-gene Primed module scored higher in Primed SMCs in every cohort (all p_adj ≤ 1.21 × 10⁻¹⁹), and each constituent gene was individually enriched, confirming out-of-sample validity (Fig. 6F). The conserved cores and bed-predominant axes are summarized in Fig. 6G.

Using counts >0 as a conservative lower bound on prevalence, *Cd34* was detected in 10.5–25.7% of Primed versus 0.1–2.1% of contractile SMCs, *Vcam1* in 21.5–51.5% versus 2.1–6.5%, and *S100b* in 0.8–6.9% versus 0.1–1.8%. *Cd34* showed the greatest specificity, with a 6-to 105-fold difference. In an independent 3-day ligation dataset^28^, Cd34⁺ cells comprised 16.1% of baseline Primed SMCs and fell to 2.7% after ligation despite expansion of the compartment. This is opposite to the expectation for de novo acquisition. Thus, *Cd34* and *Vcam1* mark substantial and specific subsets of the pre-existing pool, although single-cell dropout makes measured fractions lower bounds.

Functional enrichment across all four cohorts confirmed a contractile-low, matricellular-high architecture (Fig. 6H–J). Primed SMCs were depleted of sarcomeric machinery, vascular contraction and cGMP–PKG signalling, but enriched for ECM–receptor, proteoglycan and AGE–RAGE pathways. Migration and chemotaxis, ECM organisation and adhesion, vascular development, and skeletal/cartilage programmes were active at homeostasis. The skeletal/cartilage programme was strongest in carotid Primed SMCs, foreshadowing fibrochondrocyte fate. An integration-free metric based on cohort-level fold changes provided qualified support for a shared Primed identity: five of six Primed–Primed cohort pairs were positive, with the carotid–aortic-media comparison the exception (Supplementary Fig. 4a–c).

Partition-based graph abstraction and cross-time-point temporal flow, which do not require root selection, placed Primed SMCs as the conduit between contractile and disease-expanded states and inferred directionality from the Primed compartment despite the numerical predominance of contractile SMCs at baseline (Supplementary Fig. 5a,b). Their agreement with feeder-source and trajectory analyses identifies the homeostatic Primed pool as the dominant inferred source, while preserving the distinction between computational ancestry and direct lineage proof.

### The lesion fibromyocyte and fibrochondrocyte states are a continuation of the Primed SMC programme

Feeder and trajectory analyses identified Primed SMCs as the dominant inferred source of lesion-associated FbC. We therefore asked whether FbC retained the Primed programme or converged on an unrelated state. The Primed module (*Vcam1, Timp1, Thbs1, Cd34*) remained low in contractile SMCs at every time point in both models but was elevated in Primed SMCs, fibromyocytes and fibrochondrocytes (Fig. 7A). Module expression was highest in fibromyocytes and, although attenuated in terminal fibrochondrocytes, never returned to contractile baseline. The chondrogenic programme (*Acan, Sox9, Col2a1, Comp, Chad, Ibsp*) was restricted to fibrochondrocytes (Fig. 7B). It appeared at Week 16 in *Ldlr*^−/−^ and Week 8 in *Apoe*^−/−^ disease. *S100b* was retained across Primed SMCs and lesion-associated states (Fig. 7C). Four signatures were shared by Primed SMCs and FbC across both models and the carotid reference: the Primed surface module, a pan-vascular core, *Ly6a* and a secreted bridge programme defined by *Spp1* and *Prg4* (Fig. 7D). Fibromyocyte ECM genes and the terminal chondrogenic programme appeared only after lesion formation. FbC therefore represent progressive elaborations of a pre-existing Primed programme rather than unrelated endpoints.

**Fig. 7.**
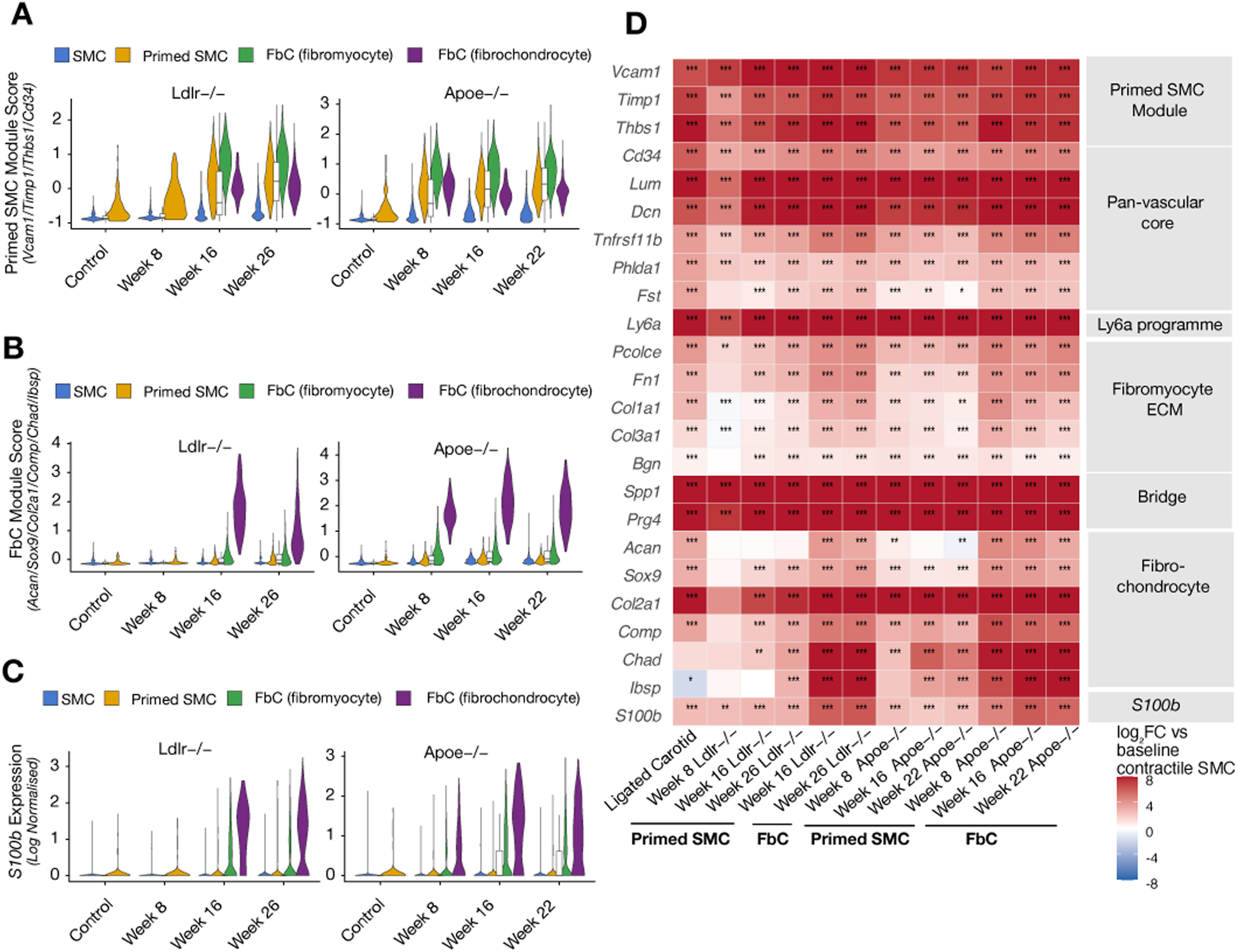
Primed identity is preserved across the Primed SMC to fibrochondrocyte trajectory. Native per-cohort annotation of contractile SMC, Primed SMC and FbC in *Ldlr*^−/−^ and *Apoe*^−/−^ disease. (A–C) Per-cell distributions across time points: Primed-module score (A), fibrochondrocyte-module score (B) and S100b expression (C). (D) Heatmap of 24 genes spanning the Primed module, pan-vascular core, Ly6a, fibromyocyte ECM, Spp1/Prg4 bridge, fibrochondrocyte programme and S100b. Colour, log₂ fold change versus each cohort’s baseline contractile SMC; stars, Wilcoxon BH-adjusted P.

### A Primed-like SMC state pre-exists in the healthy human aorta

To test for a human counterpart, we reanalysed the aortic compartment of a human vessel atlas^29^ (36,606 cells from four samples and three donors; Fig. 8A). The SMC compartment contained contractile SMC1 (2,098 cells; 99.5% MYH11⁺) and Primed-like SMC4 (3,602 cells; 97.6% MYH11⁺; Fig. 8B). Across SMCs, Primed and contractile scores anticorrelated (ρ = −0.54), and SMC4 shifted +0.84 along the Primed–contractile axis relative to SMC1 (−1.38 versus −2.22; P < 2.2 × 10⁻³⁰⁸; Fig. 8C). The Primed score localized to SMC4 (Fig. 8D), while state-resolved embedding separated SMC1, SMC4 and chondrogenic SMC5 (Fig. 8E). Versus SMC1, SMC4 upregulated VCAM1, TIMP1, CD34, LUM and TNFRSF11B (Fig. 8F). Ten of eleven core genes were concordant between human and three non-injury mouse aortic datasets (one-sided sign test P = 0.0059; Fig. 8G); THBS1 was the exception. SMC4 scored above SMC1 on the Primed module (Fig. 8H); SMC5 scored highest but had largely lost SMC identity, so Primed assignment rested on lineage retention and the Primed–contractile axis. Contractile score was highest in SMC1, retained in SMC4 and reduced in SMC5 (Fig. 8I), paralleling MYH11⁺ fractions of 99.5%, 97.6% and 34.5%. Thus, SMC4 retained SMC identity, whereas SMC5 was osteochondrogenic.

**Fig. 8.**
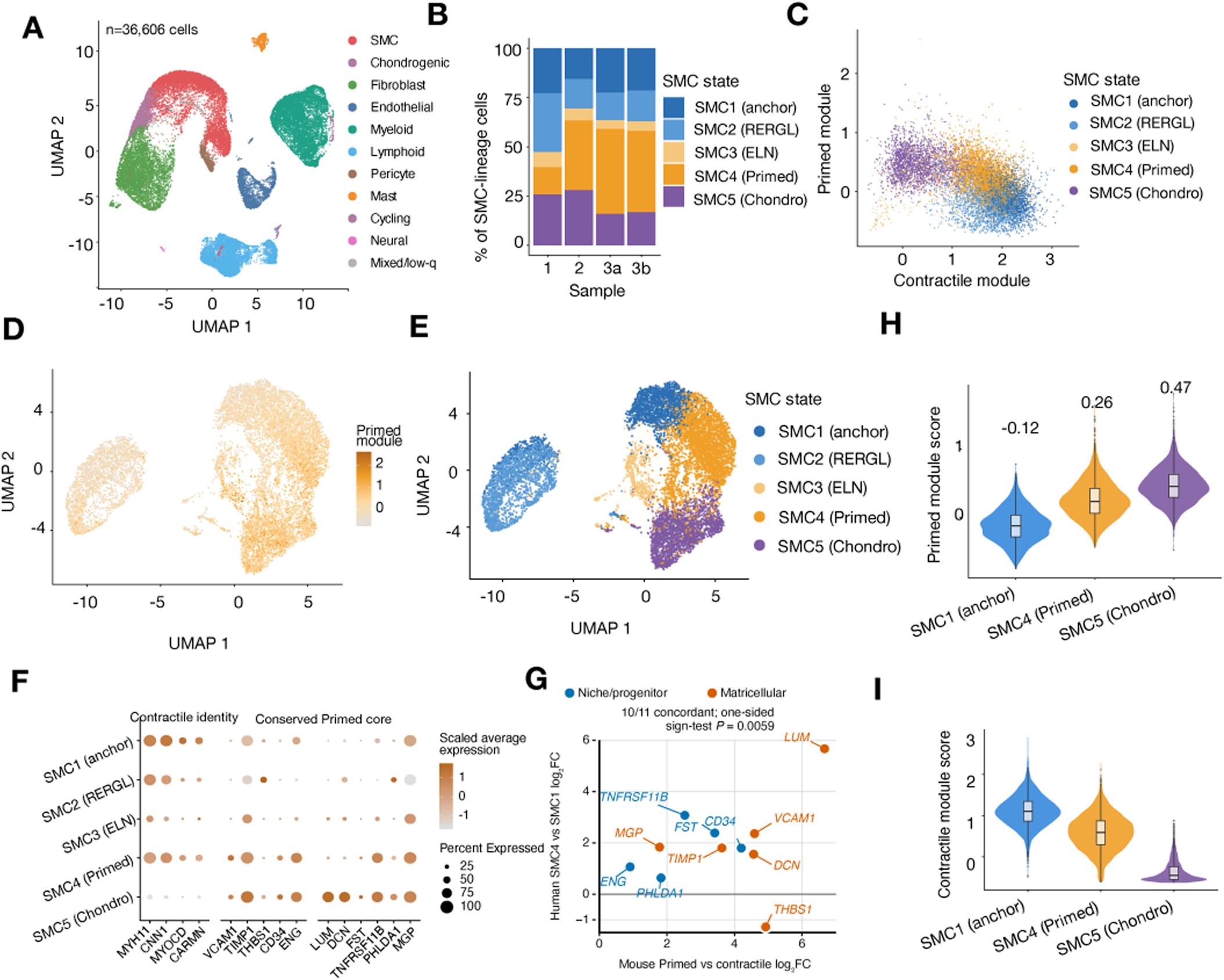
A Primed-like SMC state pre-exists in healthy human aorta. Aortic samples from a human vessel atlas^29^ processed through the mouse pipeline. (A) Integrated lineage atlas. (B) SMC-state composition across four samples. (C) Per-cell Primed-versus-contractile module score. (D) Primed-module score on the SMC embedding. (E) SMC embedding by state. (F) Lineage and conserved-core marker expression. (G) Directional concordance of eleven conserved-core genes between human SMC4 versus SMC1 and mouse Primed versus contractile SMC; ×, discordant. (H) Primed-module score for SMC1, SMC4 and SMC5. (I) Contractile-identity-module score for the same states.

## Discussion

The principal finding is that a rare homeostatic Primed SMC compartment is consistently identified as the dominant inferred ancestral source of lesion-associated SMC populations across vascular injury and atherosclerosis^1,6,7^. Convergence of independent analytical frameworks supports expansion and diversification of this pre-existing compartment rather than widespread switching of the contractile medial majority, while stopping short of direct clonal proof.

The Primed state was conserved across four baseline analyses by an integrated marker comparison and an integration-free fold-change metric, establishing that it pre-exists disease. Genetically lineage-marked aortic medial cells retained a conserved SMC core while expressing a matricellular, contractile-low and chondrogenic-poised programme consistent with later fibrochondrocyte fate. A comparable population was present in non-diseased human aorta^29^ and shifted along the same Primed–contractile axis. Ten of eleven core genes were directionally concordant across the human and three non-injury mouse aortic datasets; THBS1 was the single species-divergent exception. The state therefore precedes disease in both species, although human clonal ancestry remains unresolved.

Disease-associated contraction of the contractile compartment accompanied by Primed expansion is compatible with phenotypic switching^30,31^ and with autonomous Primed expansion alongside contractile loss. Because the models make opposing predictions about feeder identity, we tested them directly. In both atherosclerosis cohorts, the homeostatic Primed compartment was the predominant inferred source of disease-expanded Primed cells at every stage and of FbC and cycling cells at emergence. These populations subsequently became self-maintaining, while contractile SMCs were consistently depleted as feeders. The pattern persisted across neighbourhood sizes and after cell-cycle regression. Contractile substates showed internal stress-related drift without acquiring the Primed programme. Two analyses independent of root selection and pseudotime—graph abstraction and temporal flow—also placed Primed SMCs between contractile and disease-expanded states and inferred directionality from the Primed pool. Convergence across methods makes a framework-specific artefact less likely.

Previously described fibromyocytes^32^, SEM cells^25^, Lgals3⁺ pioneers^33^ and modulated or intermediate continuum states^34–36^ overlap the disease-expanded Primed compartment and share a *Vcam1, Timp1, Thbs1* core, but carry different *Ly6a, Tcf21* and *Lgals3* overlays. In many studies, transition direction was determined by a trajectory rooted in the numerically dominant contractile population, making contractile-to-modulated transitions inevitable. Across lineage-resolved cohorts, feeder-source analysis instead favoured continuity with the homeostatic Primed pool. These published states can therefore be reconciled with autonomous expansion of a pre-existing minority compartment.

Ding and colleagues assigned a Vcam1⁺ inflammatory SMC population to phenotypic switching by pseudotime^28^. Reanalysis recovered the same conserved matricellular programme but inferred a pre-existing Primed source; the original Monocle2/DDRTree direction depended on a user-specified root^37,38^. Although Vcam1 was historically considered absent from normal media and induced after lesion formation^39,40^, we detected it in baseline Primed SMCs across all cohorts. Lesional Vcam1⁺ SMCs may therefore expand from a pre-existing compartment rather than acquire Vcam1 de novo.

AdvSca1-SM progenitors, proposed to arise from mature SMCs through Klf4-dependent reprogramming^14^, share *Cd34, Ly6a* and a *Lum/Dcn/Pi16/Scara5* fibroblast signature with disease-expanded Primed SMCs. However, retained contractile identity favours a medial location for Primed SMCs. The populations may represent related components of a shared progenitor-like lineage, but spatial analysis is needed to resolve their relationship.

The strongest functional evidence for a medial Cd34⁺ progenitor comes from a dual-recombinase study of a Cd34⁺ smooth-muscle-derived transient progenitor^41^. This population generated most neointimal SMCs, and its ablation reduced hyperplasia. Although the study concluded that Cd34 was acquired after injury because baseline immunostaining was negative, its single-cell data contained a discrete Cd34⁺ subcluster at baseline (1.25% of SMCs) that expanded to 12.4% after injury. An independent 3-day ligation dataset likewise contained a baseline Cd34⁺ Primed fraction (16.1%) that fell to 2.7% despite expansion of the overall compartment^28^. *Dclk1*, the proposed mediator^41^, was enriched in baseline Primed SMCs across all four cohorts and increased further during disease. Together, these findings support constitutive Cd34 and Dclk1 expression in a pre-existing compartment rather than injury-induced acquisition by contractile SMCs.

The model also explains why Sca1-CreERT² labels few lesional SMCs^12,13^ despite recurrent Ly6a/Sca1⁺ disease populations. *Ly6a* is acquired during expansion of the homeostatic Primed pool, so pre-injury Sca1 tracing would miss much of the compartment. S100b-CreERT studies^20,21,42^ may capture the small but significant baseline *S100b*⁺ Primed fraction (∼7%); S100b knockdown also reduces neointima and a Ly6a/Sca-1⁺/α-SMA⁺ intermediate^43,44^. Intersectional Myh11 tracing gated by the more prevalent *Cd34* or *Vcam1* markers should provide broader coverage.

Therapeutically, the switching paradigm motivates suppression of modulation across contractile SMCs^30,31^. The present model instead identifies a rare Primed compartment whose expansion accompanies lesion growth. Broad anti-proliferative strategies directed at contractile SMCs may therefore affect cells that are not the main contributors to lesion-associated populations. *Cd34* and *Vcam1* enable prospective isolation for perturbation and screening, while matrix and adhesion programmes nominate candidate targets. Any intervention would require direct lineage validation and assessment of normal vascular functions.

A key limitation is that kNN feeder-source analysis, like trajectory reconstruction, infers transcriptional proximity to candidate pre-existing states rather than direct clonal ancestry. It cannot exclude convergence of contractile cells on the Primed state^38^. Nevertheless, multicolour tracing establishes oligoclonal SMC accumulation from a restricted Myh11⁺ subset^7,9,10^. Repeated recovery of the same homeostatic compartment, its dominant self-feeding, consistent downstream contributions and reciprocal contractile-source depletion make Primed SMCs the best-supported ancestral compartment currently detectable. Definitive proof requires prospective barcoding, for example by combining an SMC-specific driver with CARLIN^45^ or DARLIN^46^, and mtDNA-based clonal reconstruction in human lesions^47^.

Spatial X-inactivation analysis using a BGN 3′-UTR deletion showed that clonal intimal SMCs correspond to dominant medial clones, supporting a medial origin^48^. The same study identified a predominantly polyclonal lesion with a smaller clonal component, compatible with expansion of a pre-existing pool distributed across the media. Such expansion could appear more oligoclonal in smaller murine lesions analysed with a limited colour palette. Single-cell mtDNA genotyping has likewise identified spatially restricted and state-biased SMC expansion in human aorta^49^. Neither approach resolved the founder-cell identity. Applying mtDNA-based reconstruction within the Primed-versus-contractile framework would directly test the model in humans.

The carotid comparison comprised one pooled library per condition. It supports discovery of the Primed state, its relative representation and within-library transcriptional separation, but not between-animal inference or an estimate of absolute population expansion. A second limitation is that all mice were male: the *Myh11*-CreERT² transgene is Y-linked, and the carotid ligation cohort also used male mice only. The Primed state was also recovered in the non-sex-restricted human dataset, but female-inclusive models are required to test sex-dependent representation and responses.

In summary, carotid injury, healthy Myh11-lineage-traced aortic media and two atherosclerosis models converge on a shared architecture in which a rare pre-existing Primed SMC compartment is the dominant inferred source of lesion-associated SMC populations. The findings favour autonomous expansion and diversification of this compartment over widespread switching of the contractile medial majority. The model identifies a disease-relevant population before lesion formation, focuses mechanistic and therapeutic studies on a prospectively isolatable compartment, and defines prospective lineage tracing as the critical next test.

## Methods

### Animals and experimental models

All experimental procedures involving original mouse work for the carotid ligation cohort were approved by the University of Rochester Animal Care Committee and conformed to NIH guidelines for the care and use of laboratory animals.

### Carotid ligation model

Cdh5(PAC)-CreERT² × Ai9 tdTomato reporter mice (Taconic Biosciences #13073 × Jackson Laboratory #007909) were used to generate Cdh5–Ai9 mice. Male mice aged 6–8 weeks (∼20 g) were used; mice were housed under standard specific-pathogen-free conditions with a 12-hour light/dark cycle and food and water provided ad libitum. The Cdh5–Ai9 lineage-tracing background was part of the original experimental design but tdTomato transcripts were insufficiently detected in scRNA-seq data for reliable endothelial lineage assignment. The present study therefore defines cell identities transcriptionally.

Left common carotid artery ligation was performed as previously described^20,42^. Mice were anaesthetised with inhaled isoflurane, administered sustained-release buprenorphine (0.5-1.0 mg/kg, subcutaneous), and maintained on a temperature-controlled surgical platform. The left common carotid artery was exposed and ligated proximal to the bifurcation using 6-0 silk suture. Sham-operated mice underwent identical exposure without ligation. Arteries were harvested 14 days post-surgery for single-cell dissociation. For each condition, carotid arteries from 10 mice were pooled before dissociation and processed as one composite 10x Genomics Chromium library. The pool, rather than each mouse or vessel, was the sequencing experimental unit (n = 1 pooled library per condition). Mouse- and vessel-level identities were not retained; consequently, between-animal variability and replicate-level condition effects could not be estimated.

The sham and ligated arms analysed here form part of a wider multi-arm carotid study that additionally includes treatment arms reported separately. All arms were dissociated, sequenced, integrated and clustered together as a single object; the present study analyses only the untreated sham and ligated arms. The complete multi-arm object will be deposited on publication, with the treatment arms labelled.

Mice were euthanised by ketamine/xylazine anaesthesia followed by PBS perfusion. Common carotid arteries were dissected and immediately transferred into ice-cold FACS buffer (Thermo Fisher Scientific, Cat# 00-4222-57). Carotid artery segments from each sample were pooled and minced into fragments < 1 mm3 in enzyme digestion buffer containing Liberase TM (4 U/mL; Sigma-Aldrich, Cat# 540119001), hyaluronidase (60 U/mL; Sigma-Aldrich, Cat# H3506) and DNase I (120 U/mL; Thermo Fisher Scientific, Cat# 85836) in PBS, and incubated at 37 °C with shaking for 45–60 min. Samples were passed through a 70 µm strainer, centrifuged at 500 × g for 5 min at 4 °C, mechanically disrupted, rinsed with RPMI + 10% serum, re-pelleted and resuspended in RPMI + 10% serum. Viability was assessed by trypan blue exclusion. Dead cells were depleted using the Akadeum Microbubble Dead Cell Removal Kit. Final pellets were resuspended in RPMI + 10% serum prior to 10x Genomics library preparation.

### Healthy aortic medial SMC dataset

Healthy aortic medial SMC transcriptomes were obtained from GEO (GSE117963; GSM3316207)^10^. The present study analysed the four Myh11-CreERT²-marked (Confetti⁺) aortic medial 10x libraries (GSM3316207–GSM3316210; 12,092 lineage-positive cells). Because the Myh11-CreERT² transgene is Y-linked, all mice were male. Tamoxifen was administered intraperitoneally (1 mg in corn oil) for 10 doses beginning at 6–8 weeks of age. Cells were enzymatically dissociated, and viable Confetti-positive singlets were isolated by FACS.

### Atherosclerosis lineage-tracing cohorts

Single-cell RNA-seq datasets from *Ldlr*^−/−^ and *Apoe*^−/−^ atherosclerosis models were obtained from GEO (GSE155513)^25^. Myh11-CreERT² mice (JAX 019079) were crossed with Ai6 (RCL-ZsGreen1) reporter mice (JAX 007906) and bred onto either *Ldlr*^−/−^ (JAX 002207) or *Apoe*^−/−^ (JAX 002052) backgrounds on C57BL/6J (JAX 000664). Tamoxifen-induced recombination permanently labelled SMCs and their progeny with ZsGreen1. Because the Myh11-CreERT² transgene is Y-linked, all lineage-tracing mice were male. Mice received tamoxifen-containing chow for 2 days followed by 2 days of standard chow before initiation of a Western diet (Envigo Teklad TD.88137; 21% fat, 0.2% cholesterol) at 8 weeks of age. *Ldlr*^−/−^ mice were analysed at 0, 8, 16 and 26 weeks of diet, and *Apoe*^−/−^ mice at 0, 8, 16 and 22 weeks.

The *Ldlr*^−/−^ and *Apoe*^−/−^ cohorts were analysed as separate objects, each independently Harmony-integrated, embedded and clustered together with its own Western-diet timepoints, with no joint cross-cohort integration. The two cohorts share a single Control (pre-diet) timepoint rather than each carrying an independent Control: the same 6,562 Myh11-CreERT² Control cells (3,506 FACS-negative and 3,056 ZsGreen⁺) were included in, and independently re-embedded and re-clustered within, each cohort object. Because every Control cell is common to both objects, the per-cohort Control Primed SMC counts (*Ldlr*^−/−^ n = 141; *Apoe*^−/−^ n = 100) reflect independent clustering and lineage-gating of one shared baseline rather than two biological samples. The Western-diet timepoints are cohort-specific. Aortic tissue dissection was restricted to the ascending aorta, brachiocephalic artery and descending thoracic aorta. The aortic medial dataset sampled adventitia-free whole aortae^10^.

### scRNA-seq library preparation and alignment

#### Carotid ligation dataset

Pooled carotid arteries were processed using the 10x Genomics Chromium Single Cell 3′ v3 platform and sequenced on an Illumina platform at the University of Rochester Genomics Research Centre. BCL files were converted to FASTQ using Cell Ranger v7.0.1 mkfastq and aligned with Cell Ranger count to the mm10 reference (refdata-gex-mm10-2020-A) with the tdTomato reporter sequence appended (refdata-gex-mm10-2020-A-tdTomato). Cell Ranger recovered 2,285 sham and 4,925 ligated cells; median per-cell metrics were 4,569 genes and 18,643 UMIs (sham) and 3,378 genes and 10,989 UMIs (ligated). After quality control and doublet removal, 1,973 sham and 4,272 ligated cells were retained for downstream analysis (6,245 cells analysed; sham and ligated arms of the 26,426-cell multi-arm object).

#### Public datasets

Raw FASTQ files from public datasets were downloaded from the SRA and processed using a unified pseudoalignment workflow with kallisto v0.51.1, bustools v0.44.1 and kb-python v0.29.1 (10x v2/v3 workflow^50,51^). All datasets were aligned to the 10x Genomics mm10 2020-A reference. For the atherosclerosis cohorts, the ZsGreen1-WPRE reporter sequence was appended to the reference to enable simultaneous quantification of endogenous transcripts and lineage reporter expression. Filtered count matrices were generated in h5ad format and imported into Seurat v5.5.0 using zellkonverter^52^. Ensembl identifiers were converted to mouse gene symbols using org.Mm.eg.db. for downstream variable-feature analyses. The human aortic atlas was derived from the original Cell Ranger alignment^29^. Although Cell Ranger was used for the in-house carotid and human aortic datasets and kallisto/bustools for public datasets, both approaches yield comparable gene-level quantification.

Independent acute-injury dataset and marker-capture analysis. An independent 3-day carotid-ligation dataset (sham and ligated; SRA PRJNA628067)^28^ was downloaded and processed through the same pseudoalignment, normalisation, Harmony integration and clustering pipeline described above; the Primed and contractile SMC compartments were identified by the conserved Primed module (*Vcam1, Timp1, Thbs1, Cd34*) and the contractile module (*Myh11, Myocd, Cnn1, Carm*n), respectively. Marker-capture analysis was then applied uniformly across all five murine datasets (carotid ligation, aortic media, *Ldlr*^−/−^, *Apoe*^−/−^ and the 3-day ligation cohort). Within the baseline (sham/control) Primed and contractile compartments — restricted, for the lineage-traced atherosclerosis cohorts, to ZsGreen1-WPRE–positive cells — the fraction of cells expressing each candidate marker (*Cd34, Vcam1, S100b*) was computed as the proportion with detected transcript (counts > 0), an intentionally conservative lower bound on promoter activity given single-cell dropout. Percentages are reported with Wilson 95% confidence intervals, and Primed-versus-contractile enrichment as the ratio of these fractions. Reported single-cell percentages from the smooth-muscle–derived transient progenitor cell study^41^ were taken from that study’s published single-cell re-clustering of the SMC compartment; those data were not reprocessed here.

### Quality control and filtering thresholds

Quality control was performed independently for each cohort, with parameters chosen according to the cell-loss biology of each preparation. For all four cohorts, cells were filtered using nFeature_RNA and nCount_RNA outlier thresholds defined per sample as the 75th percentile + 1.5 × IQR, requiring a minimum of 200 detected genes per cell and a minimum of 3 cells per gene. Doublets were identified and removed using DoubletFinder (v2.0.6)^53^ with an expected doublet rate of 5%, pN = 0.25 and pK selected by parameter sweep maximising the mean-variance-normalised bimodality coefficient. Mitochondrial transcript fraction was quantified against the mouse mitochondrial gene set and applied uniformly as an exclusion criterion across all four cohorts at 10%. Cells exceeding 10% mitochondrial content were removed as likely dissociation-damaged. Mitochondrial transcript fractions were generally low across the carotid atlas after QC, with most clusters concentrated below the 10% exclusion threshold. Elevated mitochondrial content was restricted to a small number of discrete clusters rather than broadly affecting the major contractile or Primed SMC compartments, supporting recovery of a high-complexity Primed state after QC. This QC check does not establish absolute cell abundance or expansion. As a robustness check, applying a stricter ≥450-gene threshold removed none of the contractile or Primed SMCs analysed, all of which were high-complexity cells (per-cohort median 1,357–3,634 genes), leaving the Primed compartment, its module separation and *Cd34/Vcam1* enrichment unchanged.

### SMC lineage assignment

In the healthy aortic medial dataset, all analysed cells had been pre-enriched for *Myh11* lineage identity by FACS in the original study and were treated as lineage-positive without further filtering; in healthy aorta, cell death and phagocytic clearance are minimal, so reporter enrichment is not appreciably confounded by protein transfer. For the *Ldlr*^−/−^ and *Apoe*^−/−^ cohorts, FACS-sorted ZsGreen1-positive and ZsGreen1-negative fractions were merged prior to analysis and lineage was assigned by reporter transcript rather than by the sort gate. FACS reports ZsGreen protein, which in atherosclerotic plaques can be acquired both through Cre-mediated recombination and through phagocytic engulfment of ZsGreen1-positive cells and debris. Notably, cells discordant for the sort gate and the transcript (FACS-positive, transcript-negative) were predominantly myeloid (66–69% myeloid-marker⁺; *Cd68, Adgre1, Lyz2, Csf1r, C1qa*), consistent with phagocytic acquisition of reporter protein rather than heritable labelling, with a minority (25–28%) of SMC-marker⁺ cells showing genuine transcript dropout (Supplementary Fig. 6a–f). Defining lineage by the integrated transgene transcript excludes these false-positives and recovers genuinely recombined cells mis-assigned to the negative sort gate.

Lineage-positive cells were defined by ZsGreen1-WPRE expression exceeding 3.17 log-normalised counts. This threshold was set by maximising Youden’s J against the FACS sort gate in the *Ldlr*^−/−^ reference cohort (n = 26,673 cells; AUC = 0.90, sensitivity = 0.76, specificity = 0.97) and applied globally to both atherosclerosis cohorts and across all timepoints (Supplementary Fig. 7a–f). FACS-positive and FACS-negative cells were clustered jointly (Harmony integration over sample) to provide a common transcriptional reference, and lineage status was assigned within that shared embedding. Because this criterion requires detectable transgene transcript, it is deliberately conservative.

### Normalisation, integration and clustering

Count matrices were normalised using Seurat NormalizeData (scale factor = 10,000). Highly variable genes were identified with FindVariableFeatures (3,000 genes; VST), followed by ScaleData, and PCA was computed on the 3,000 variable features (50 PCs). Within each dataset, libraries (and timepoints, for the atherosclerosis cohorts) were integrated using Harmony v1.2.3^54^. Harmony used sample and timepoint metadata for the atherosclerosis cohorts and sample alone for the carotid and aortic medial datasets. UMAP embeddings and nearest-neighbour graphs were generated from Harmony-derived dimensions 1–41 (carotid), 1–45 (aortic medial), and 1–30 (*Ldlr*^−/−^ and *Apoe*^−/−^). Clustering used FindClusters at resolution 0.4 for all murine cohorts, chosen to resolve the principal mural, adventitial, endothelial and immune compartments, including the contractile SMC and Primed SMC compartments at single-population granularity, without fragmenting into sub-clusters lacking distinct marker profiles. At the clustering resolution used (0.4), fibromyocytes and fibrochondrocytes formed a single FbC cluster in the *Ldlr*^−/−^ dataset and two adjacent clusters in the *Apoe*^−/−^ dataset; for cross-model comparability the two *Apoe*^−/−^ clusters were treated as a single FbC compartment in all composition, feeder and trajectory analyses. Each dataset was analysed independently; no joint cross-dataset Harmony integration or co-clustering was performed. Human aortic data were processed in two stages. All vessel-wall cells were normalised, scaled, and integrated across donors with Harmony, with UMAP embedding and clustering on Harmony dimensions 1–30 (resolution 0.5). Vascular SMCs were then subset and independently re-embedded and re-clustered across donors on Harmony dimensions 1–20 (resolution 0.3).

### Differential expression and module scoring

Differential expression was performed with Seurat FindMarkers using two-sided Wilcoxon rank-sum tests. min.pct = 0.1 set the prevalence floor for a gene to be tested. Unless otherwise stated, logfc.threshold = 0.25 was applied during testing. For Fig. 1C, FindMarkers was run with logfc.threshold = 0, and significance was assigned post hoc at |average log₂FC| > 0.25 and Bonferroni-adjusted P < 0.05. Cross-cohort tier and enrichment analyses were likewise run without a log-fold-change filter, with the effect-size cutoff applied post hoc. Within-cohort gene-level P values used Seurat’s default Bonferroni correction across the full assayed feature set; cross-cohort comparison families used Benjamini-Hochberg correction. Figure 1G reports the Bonferroni-adjusted values from the Fig. 1C differential-expression analysis. For the descriptive Fig. 1D display, the 15 Primed-enriched genes were selected by decreasing average log₂FC, whereas the 15 contractile-SMC-enriched genes were selected by smallest Bonferroni-adjusted P; bar lengths show average log₂FC. Module scores were calculated with AddModuleScore using the gene sets listed in Supplementary Table 4. In Extended Data Fig. 9, a five-gene contractile core (Myh11, Acta2, Cnn1, Tagln, Myocd) was scored alongside the progenitor-like and matricellular modules. Human scores used a five-gene Primed set (VCAM1, TIMP1, THBS1, CD34, ENG), an eight-gene contractile set for the Primed–contractile axis (Fig. 8C) and a four-gene contractile-identity set (MYH11, CNN1, MYOCD, CARMN) for state comparisons (Fig. 8I). The Primed SMC module comprised *Vcam1, Timp1, Thbs1* and *Cd34*; the strict contractile SMC module comprised *Myh11, Cnn1, Myocd* and *Carmn*; and the fibrochondrocyte module comprised *Acan, Sox9, Col2a1, Comp, Chad* and *Ibsp*. Figure-specific extended panels were applied where indicated. Figure 1E compared the Primed and contractile scores separately using two-sided cell-level Wilcoxon rank-sum tests; the two displayed P values are raw and were not adjusted across the pair of tests. Other module-score comparison families were BH-adjusted unless otherwise stated. All Fig. 1C,E,G tests compare captured cells within the single pooled sham library and therefore quantify within-library state separation rather than between-animal variation.

Cross-cohort signature architecture: Conserved Primed SMC signatures were identified by intersecting significantly up-regulated genes across all four cohorts (BH-adjusted p < 0.05, log2FC > 0.25, expressed in ≥ 10% of Primed SMC cells per cohort). Genes meeting cross-cohort criteria were stratified into pan-vascular conserved cores versus bed-specific axes by cohort-restricted enrichment patterns.

Cross-cohort Primed SMC versus contractile SMC differential expression was performed on n = 101 (sham carotid), n = 240 (aortic medial), n = 141 (*Ldlr*^−/−^ Control) and n = 100 (*Apoe*^−/−^ Control) Primed SMC; lineage-positive cells were defined by ZsGreen1-WPRE > 3.17. Because the *Ldlr*^−/−^ and *Apoe*^−/−^ cohorts share a single Control timepoint, the *Ldlr*^−/−^-Control and *Apoe*^−/−^-Control comparisons are the same baseline cells re-clustered within each cohort object rather than independent samples, so the four comparisons span three independent biological baselines. Significance was assessed by two-sided Wilcoxon testing with Benjamini–Hochberg correction across the comparison family.

Cross-cohort cluster similarity. ClusterFoldSimilarity (CFS v1.8.0)^55^ was applied to the QC-filtered, independently log-normalised cohort objects, restricted to baseline/uninjured cells (sham carotid; control *Ldlr*^−/−^/*Apoe*^−/−^; all healthy aortic medial), lineage-gated where applicable (ZsGreen>3.17 for *Ldlr*^−/−^/*Apoe*^−/−^; Confetti-sorted medial cells for the aorta), and to the shared mesenchymal cell-type roster using final cohort-specific cluster annotations. Although clusters were defined on Harmony-integrated embeddings, similarity scores were computed solely from shared-gene fold-change signatures of the uncorrected RNA assay (13,830 common features; topNFeatures = 5, nSubsampling = 30) and did not use Harmony embeddings or batch-corrected expression. Compositions were compared at native depth and after downsampling to 120 cells per cell type (minimum 12).

### Functional enrichment analyses

Differentially expressed genes from each Primed SMC versus contractile SMC comparison were partitioned into upregulated (BH-adjusted p < 0.05, log2FC > 0.25) and downregulated (BH-adjusted p < 0.05, log2FC < −0.25) lists. Over-representation analysis (ORA) was performed independently on each list using clusterProfiler v4.16.0^56^: enrichGO (OrgDb = org.Mm.eg.db v3.22.0^57^) for Biological Process and Cellular Component ontologies, and enrichKEGG (organism = “mmu”). Mouse gene symbols were converted to Entrez identifiers with AnnotationDbi::select against org.Mm.eg.db; unmapped symbols (Ensembl IDs, RIKEN cDNAs, predicted genes) were excluded, and the mapped DE-tested gene set served as the per-cohort background universe. Significance thresholds were pvalueCutoff = 0.05 and qvalueCutoff = 0.2 with Benjamini–Hochberg correction^58^. Terms reaching BH-adjusted p < 0.05 in at least one cohort were retained for cross-cohort visualisation; near-redundant terms with substantially overlapping gene sets were de-duplicated to the canonical term. In the cross-cohort dot plots, dot size encodes gene ratio and dot fill encodes signed −log10(BH-adjusted p), the sign reflecting enrichment in the upregulated (positive) or downregulated (negative) list.

All four cohorts (carotid, aortic medial, *Ldlr*^−/−^ and *Apoe*^−/−^) were analysed by ORA. The cross-cohort GO Cellular Component and KEGG dot-plots include all four cohorts. The GO Biological Process dot-plot is restricted to the three cohorts with a disease arm (carotid, *Ldlr*^−/−^ and *Apoe*^−/−^).

### Cell-origin and trajectory analyses

kNN feeder-source analyses were performed in 20-dimensional Harmony space (a lower-dimensional space than clustering, to stabilise the neighbour-distance metric); UMAP coordinates were used only for visualisation.

#### kNN feeder-source statistical framework

To test whether disease-associated populations arise predominantly from autonomous expansion of pre-existing subpopulations rather than de novo phenotypic switching, a unified kNN feeder-source framework was applied to the carotid ligation, *Ldlr*^−/−^ and *Apoe*^−/−^ datasets. For each query cell, the k = 15 nearest neighbours were identified among earlier-condition reference cells (sham for the carotid cohort; preceding timepoints for the atherosclerosis cohorts) in 20-dimensional Harmony space, and source-cluster contributions (“feeder fractions”) were compared with expected reference-pool proportions as an observed/expected enrichment. Bootstrap confidence intervals were computed from 50 bootstrap iterations and enrichment p-values from 500 permutations of the reference-cluster labels (one-sided, Laplace-corrected), with Bonferroni correction across reference clusters within each query. Reference sets were chosen per question: SMC-lineage queries (Primed SMC, Cycling Primed SMC, AdvFb1) used a mesenchymal-restricted reference (contractile SMC, Primed SMC, AdvFb1, AdvFb2, pericyte, mural), whereas the Mac negative control used the full reference. Multi-k sweeps (k = 5–50) were performed as sensitivity analyses. Small query sets (e.g. Cycling Primed SMC, n ≈ 20) are reported with direction and confidence intervals and treated as suggestive; sets of fewer than five cells were not tested. This framework tests transcriptional ancestry as the closest pre-injury or pre-disease state to each query cell, rather than clonal lineage.

Cycling Primed SMC were defined as ligated Primed SMC that were both in S/G2M phase (Seurat CellCycleScoring with the cc.genes.updated.2019 sets, title-cased for mouse) and expressed each of Mki67, Top2a and Birc5 (counts > 0); this phase-and-marker intersection (∼20 cells) avoids the over-calling of phase alone.

#### Trajectory inference

Trajectory topology was inferred with Slingshot using dual rooting as a falsification test^59^. Each lineage was computed with the root placed alternately at the contractile/SMC cluster and at the Primed SMC cluster. Slingshot establishes connectivity and topology only; lineage directionality is adjudicated by the kNN feeder analysis, not by the trajectory, and pseudotime is presented as a topological ordering rather than a direct lineage trace. Lineage inference (getLineages) and pseudotime (getCurves) were computed in Harmony space — 20 dimensions on the carotid mesenchymal compartment, and 10 dimensions on the *Ldlr*^−/−^ and *Apoe*^−/−^ SMC-fate compartments (SMC/Primed SMC/FbC/Cycling); curves were embedded to UMAP for display. For the carotid cohort, a cell-cycle-regressed re-integration (regressing S and G2M scores, re-running Harmony) confirmed that the topology is invariant to cell-cycle structure.

#### Independent reclustering validation

To validate the Primed SMC identity independently of the integrated sham–ligated atlas, sham carotid cells passing QC (1,973 cells) were reclustered separately without integration (independent PCA, neighbour graph and UMAP). Seven clusters were identified at resolution 0.4 and assessed for Primed SMC marker expression using AddModuleScore; reciprocal barcode mapping between the sham-only clustering and the integrated-atlas annotation confirmed the Primed SMC identity independently of the joint embedding.

Mitochondrial DNA (MtDNA) variant analysis. Mitochondrial genotypes were called from the carotid ligation library using the alignment generated at sequencing and processed with mgatk. Per-cell, per-position allele counts were imported with ReadMGATK (Signac v1.17.1; Seurat v5.5.0) cells with mean mtDNA coverage ≥ 5 were retained (2,227 cells), of which 1,300 mapped to annotated ligated populations (SMC, Primed SMC, Cycling Primed SMC, Cycling Myeloid, AdvFb). Candidate substitutions were enumerated with IdentifyVariants across the mouse mitochondrial genome (all 48,897 possible substitutions; 3 alternate alleles × 16,299 bp) and filtered under a high-confidence setting (detected in ≥ 5 cells, strand correlation ≥ 0.65, VMR > 0.01; the standard Signac criteria) and a relaxed exploratory setting (≥ 3 cells, strand correlation ≥ 0.50, VMR > 0.005). No variant passed either filter: a single site was confidently detected in ≥ 3 cells and it fell below the dispersion threshold (VMR ≤ 0.005), so no site satisfied both the detection and dispersion criteria and the strand-correlation requirement never became the deciding factor. Mean alternative-allele frequencies sat at the noise floor (≈ 10⁻⁴) despite adequate per-position coverage (up to ∼150×), with strand correlation undefined for nearly all sites as expected for 3′ GEX data. No heteroplasmy-based clonal reconstruction or population-enrichment testing (positivity threshold 10% heteroplasmy) was therefore performed.

To assess whether the inferred SMC-fate architecture was specific to the kNN-feeder or Slingshot methods, the SMC-fate clusters (contractile, Primed, and where present fibromyocyte, fibrochondrocyte and cycling) were re-embedded independently (PCA followed by Harmony integration on sample) and analysed by two orthogonal approaches. For partition-based graph abstraction, a connectivity value was computed for each cluster pair as the ratio of observed to expected edges in the k-nearest-neighbour graph, with expected edges derived from a degree-preserving null model that controls for cluster size^60^. For temporal-flow analysis, the k = 30 nearest earlier-stage neighbours of every later-stage cell were identified in the shared embedding for each adjacent timepoint pair, and the cluster identity of those neighbours tabulated; enrichment was expressed as observed versus expected neighbour composition given the earlier-stage cluster frequencies. This cross-timepoint neighbour analysis is conceptually related to optimal-transport ancestor–descendant inference^61^.

### Statistical analysis

Analyses were performed in R v4.6.0 on macOS Apple Silicon (aarch64-apple-darwin23), with Seurat v5.5.0. Exact P values are reported with the adjustment method stated. Values below the double-precision limit are reported as P < 2.2 × 10⁻³⁰⁸. Where R returned P = 0 from the large-sample Wilcoxon approximation, P was recomputed on the log scale from the same tie-corrected statistic (Fig. 2D; Extended Data Fig. 4j). Permutation feeder P values are Laplace-corrected and floored at the permutation resolution ((1/501) × n_ref).

Within-cohort differential-expression P values were Bonferroni corrected; cross-cohort differential-expression and functional-enrichment P values were Benjamini-Hochberg corrected. Two-sided Wilcoxon rank-sum tests were used for exploratory comparisons of cell-level distributions. Cells are observations nested within libraries rather than biological replicates; in pooled datasets, these tests quantify within-library separation and do not estimate between-animal variability. Feeder bootstrap analyses used 50 iterations and permutation tests used 500 iterations, with Laplace-corrected empirical P values. Cluster proportions are reported as percentages of total recovered cells for the carotid cohort or lineage-positive cells for the atherosclerosis cohorts. For the carotid comparison, 10 vessels per condition were pooled into one library, giving n = 1 pooled library per condition. Sham-versus-ligated composition and recovered-cell counts are therefore descriptive; no replicate-level hypothesis test was performed, and these measurements do not by themselves estimate absolute tissue abundance. The *Ldlr*^−/−^ and *Apoe*^−/−^ timepoints were also pooled without biological replicates, and the two cohort objects share one baseline Control. Concordant recovery of the Primed state in healthy aortic media, *Ldlr*^−/−^ and *Apoe*^−/−^ datasets was treated as cross-dataset corroboration of state identity, not as biological-replicate-level statistical power for the pooled carotid comparison. No statistical methods were used to predetermine sample size. Group allocation for surgery was not randomised and surgical procedures were not blinded, owing to the clear anatomical distinction; downstream computational analyses were not formally blinded.

## Data availability

Original carotid ligation scRNA-seq data have been deposited in GEO under accession [GSE to be assigned]. The complete multi-arm carotid object (26,426 cells) is deposited with the treatment arms labelled Treatment X and Treatment Y; the underlying treatment-arm identities are released on publication of the intervention study. Public datasets reanalysed here are available from GEO accessions GSE117963 (healthy medial SMCs; GSM3316207–GSM3316210) and GSE155513 (*Ldlr*^−/−^ and *Apoe*^−/−^ cohorts). The independent 3-day carotid-ligation dataset is available from the SRA under accession PRJNA628067. The nondiseased human aortic data were obtained from the processed single-cell dataset of human cardiac arteries deposited by Hu et al. on Zenodo (https://zenodo.org/records/6032099).

## Code availability

All analysis code used to generate the main, Extended Data and Supplementary figure panels, the Supplementary Tables and the analyses reported in the text, including the kNN feeder-source framework, the Slingshot trajectory analyses and the marker-capture analysis, has been deposited in Zenodo under an MIT licence, with a README and a panel-to-script map. DOI: [to be provided upon submission].

## Acknowledgements

The authors acknowledge the contribution of the Dublin City University, School of Biotechnology project research funding programme to T-S.L.

## Author contributions

S.W., S.A., T-S.L., I.K., K.U., M.K., and P.A.C. performed all the computational analysis. W.L. performed the surgical carotid-ligation. E.M.R., A.F., and P.A.C. designed the experiments, wrote the manuscript and edited it for submission.

## Extended Data Figures

**Extended Data Fig. 1.**
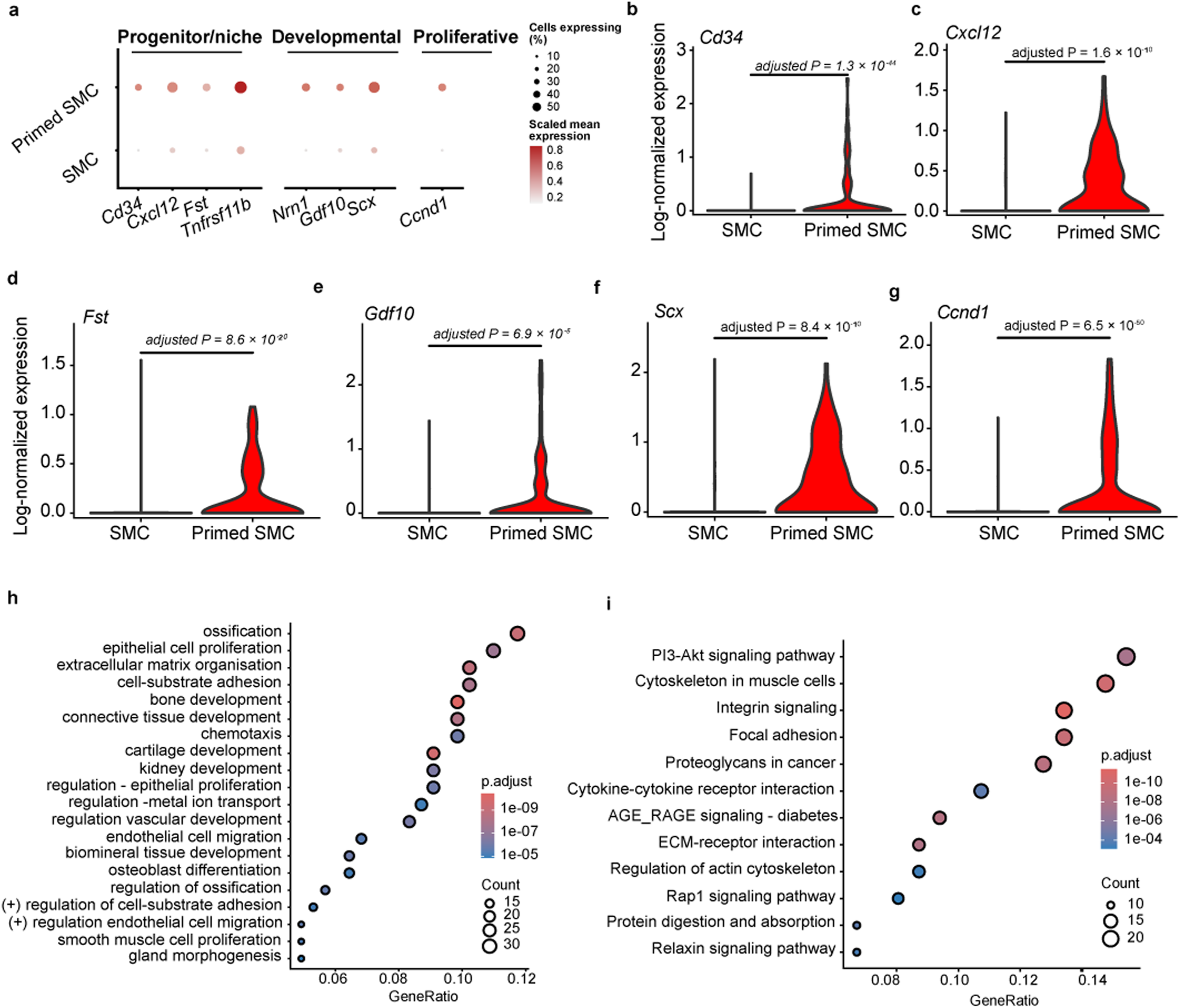
The Primed SMC progenitor-associated programme is constitutively present in the uninjured artery. (a) Dot plot of the Primed programme grouped as progenitor/niche (*Cd34, Cxcl12, Fst, Tnfrsf11b*), developmental (*Nrn1, Gdf10, Scx*) and proliferative (*Ccnd1*) genes (dot size, percentage expressing; colour, scaled average expression). (b-g) Violins in contractile versus Primed SMCs: *Cd34* (Bonferroni-adjusted P = 1.3 × 10⁻⁴⁴), *Cxcl12* (1.6 × 10⁻¹⁰), *Fst* (8.6 × 10⁻²⁰), *Gdf10* (6.9 × 10⁻⁵), *Scx* (8.4 × 10⁻¹⁰) and *Ccnd1* (6.5 × 10⁻⁵⁰) (two-sided cell-level Wilcoxon rank-sum tests; Seurat Bonferroni correction). (h,i) Over-representation analysis of genes up-regulated in Primed SMCs: top GO Biological Process terms (h) and KEGG pathways (i) (dot size, gene count; colour, adjusted P; x-axis, gene ratio).

**Extended Data Fig. 2.**
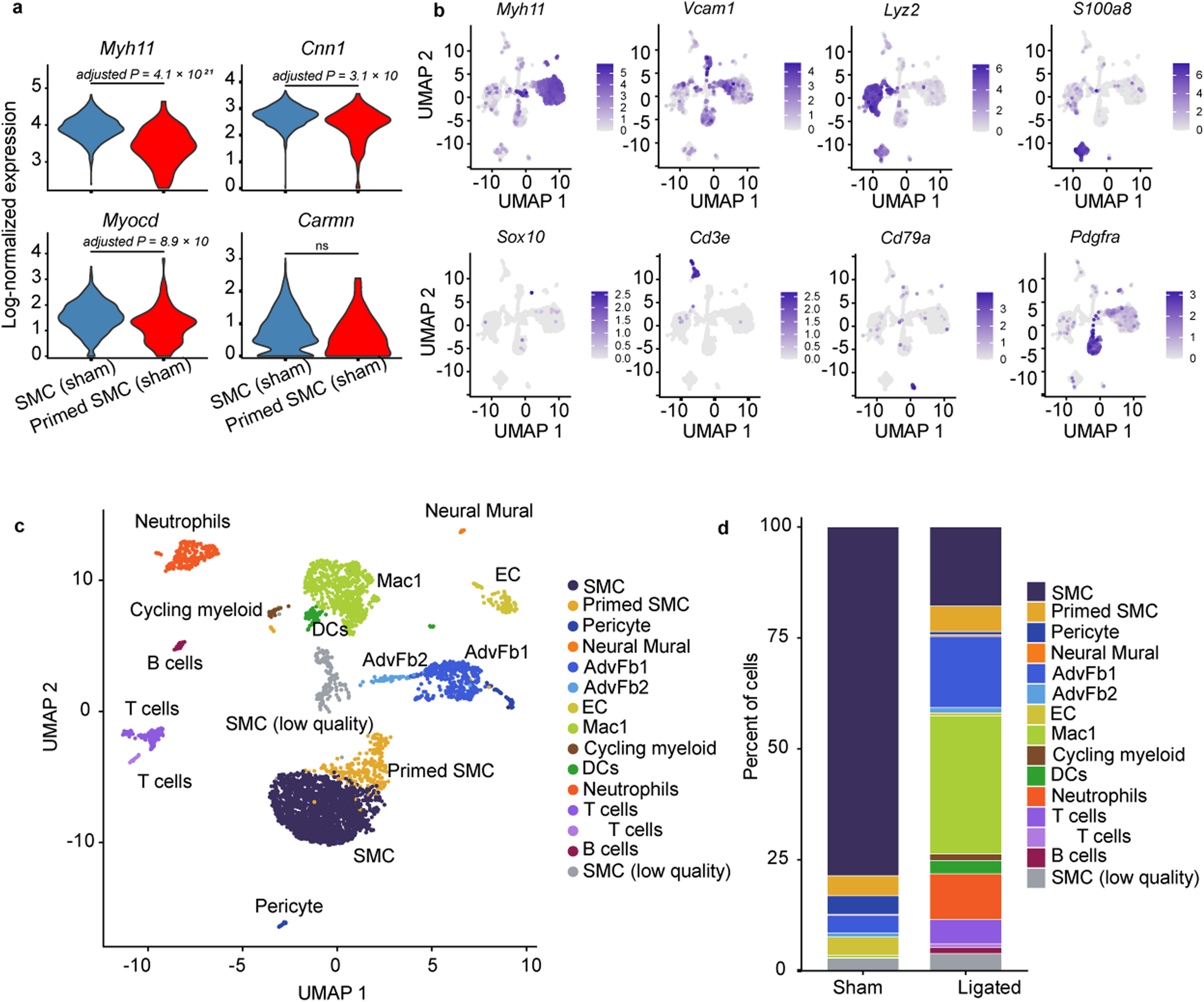
Whole-atlas markers and balanced down-sampling of sham versus ligated carotid scRNA-seq. (a) Violins of canonical contractile markers (*Myh11, Cnn1, Myocd, Carmn*) in sham SMC versus Primed SMC — retained but attenuated in Primed SMC (Wilcoxon two-sided). (b) Feature plots of compartment markers: *Myh11* (SMC), *Vcam*1 (Primed SMC), *Lyz2* and *S100a8* (innate immune), *Sox10* (neural mural/Schwann), *Cd3e* (T cell), *Cd79a* (B cell), *Pdgfra* (fibroblast). (c) UMAP after balanced down-sampling equalising Sham and Ligated cluster representation, showing that the Primed cluster remains represented after balancing cell numbers. This sensitivity analysis addresses sampling imbalance but does not establish absolute expansion. (d) Composition after balanced down-sampling.

**Extended Data Fig. 3.**
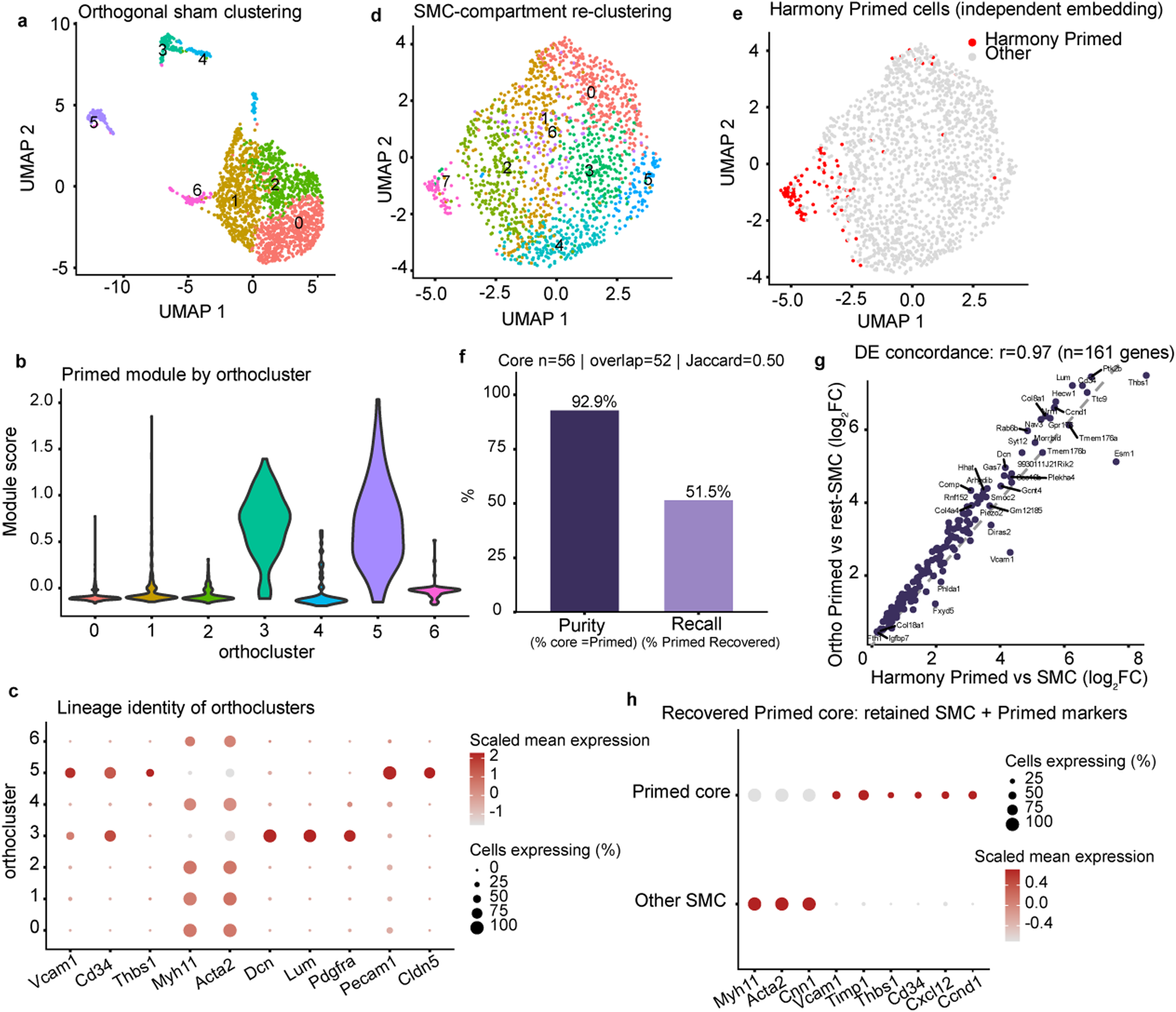
An independent, integration-free re-clustering recovers the Primed SMC pool. (a) Orthogonal re-clustering of sham cells using a separate PCA, graph and UMAP, resolving seven clusters (orthoclusters 0–6). (b,c) Primed module score (b) and lineage-marker dot plot (c) across orthoclusters: the two module-high whole-sham clusters are adventitial fibroblast (Dcn, Lum, Pdgfra) and endothelial (Pecam1, Cldn5), not SMC — the module cross-reacts with these lineages. (d,e) Re-clustering restricted to the SMC compartment (d), with integrated (Harmony) Primed SMC overlaid in red (e); these cells occupy a coherent region of the independent embedding. (f) A single SMC sub-cluster (the recovered Primed core, n = 56) is 92.9% Harmony Primed and recovers 51.5% of the Primed pool (overlap 52/101; Jaccard 0.50). (g) DE concordance: log₂FC of the orthogonal Primed-versus-rest-SMC contrast against the integrated Primed-versus-SMC contrast for shared significant genes (Pearson r = 0.97; n = 161). (h) The recovered core retains contractile identity (*Myh11, Acta2, Cnn*1) while expressing Primed markers (*Vcam1, Timp1, Thbs1, Cd34, Cxcl12, Ccnd1*).

**Extended Data Fig. 4.**
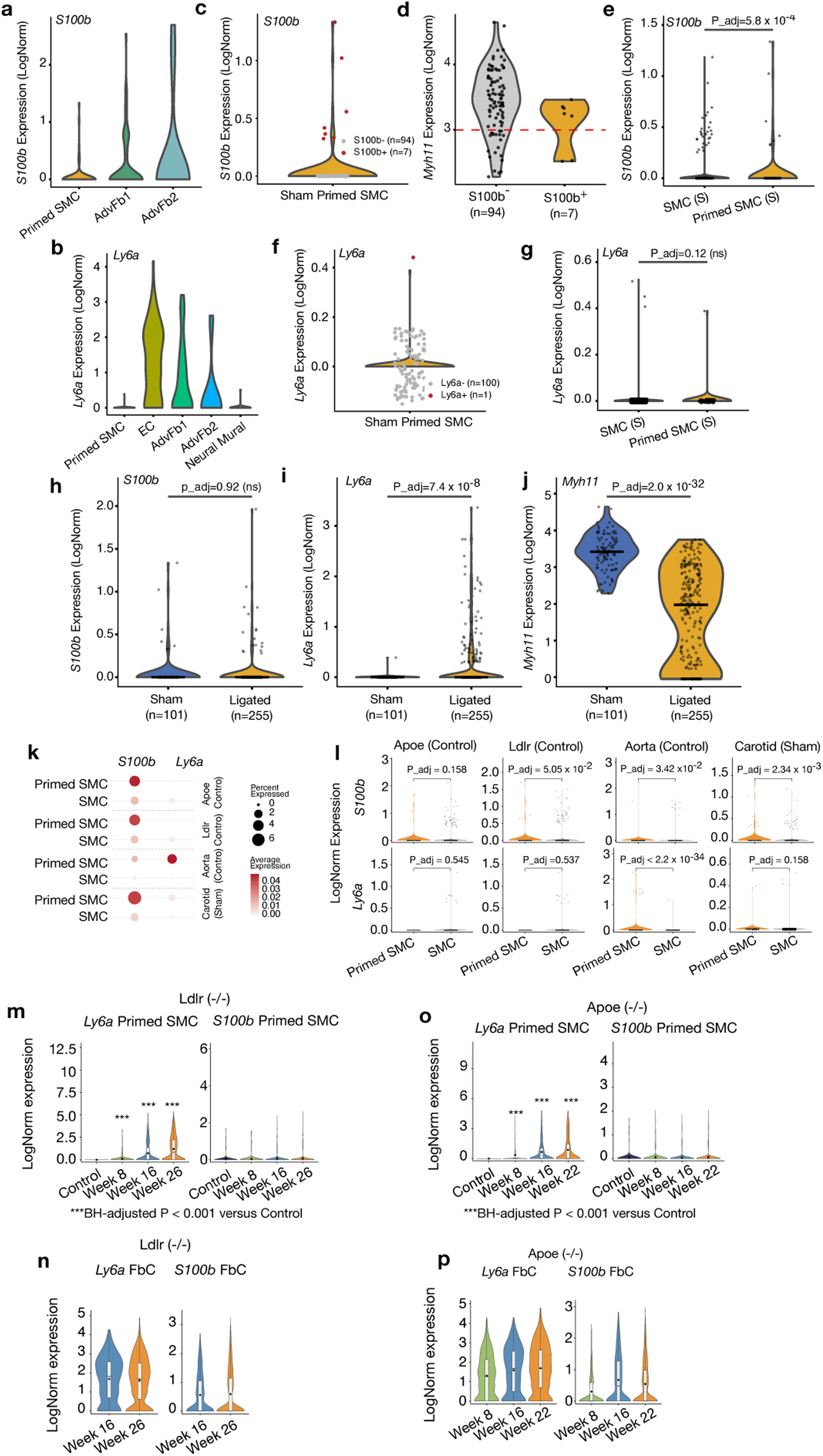
S100b and Ly6a mark complementary Primed programmes at homeostasis and during disease. (a–j) Carotid homeostasis and ligation. (a) S100b and (b) Ly6a across sham vascular populations. (c) S100b⁺ sham Primed SMC. (d) Myh11 in S100b⁺ versus S100b⁻ Primed SMC. (e) S100b enrichment in sham Primed versus contractile SMC. (f,g) Ly6a detection and comparison in sham Primed SMC. (h–j) S100b, Ly6a and Myh11 in sham versus ligated Primed SMC. (k–p) Cross-cohort baseline and disease progression. (k) S100b and Ly6a dot plot in Primed versus contractile SMC at baseline across four cohorts. (l) Corresponding violins. (m,n) Ly6a and S100b in *Ldlr*^−/−^ Primed SMC (m) and FbC (n) across available time points. (o,p) Corresponding *Apoe*^−/−^ analyses. Wilcoxon tests with BH correction; small sham-only subsets are descriptive.

**Extended Data Fig. 5.**
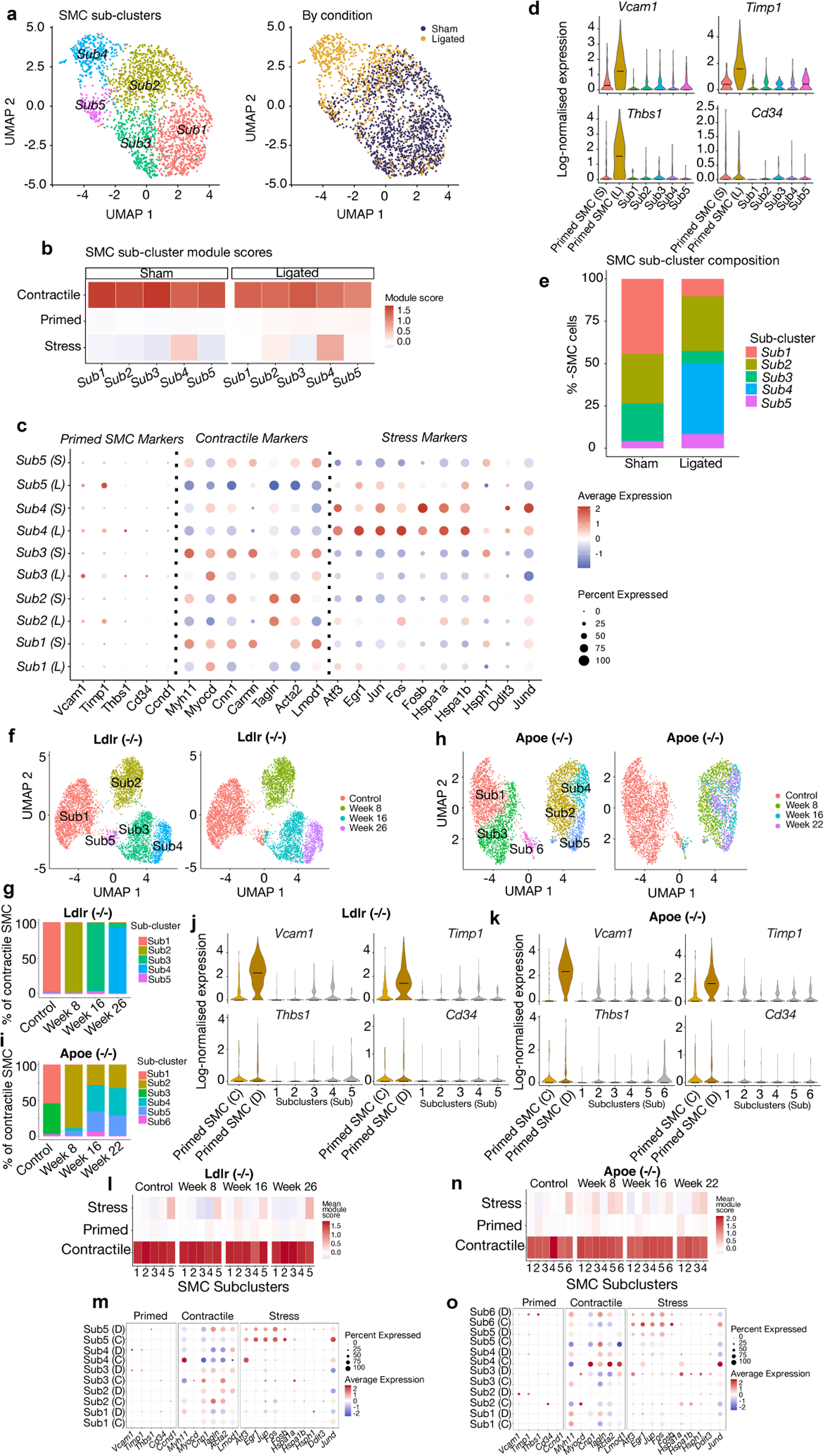
Contractile SMC subclusters do not acquire the Primed programme after injury or during atherosclerosis. (a–e) Carotid sham and ligation. (a) Contractile subclusters by condition. (b) Contractile, Primed and stress module scores. (c) Marker dot plot. (d) Vcam1, Timp1, Thbs1 and Cd34 across Primed SMC and contractile subclusters. (e) Subcluster composition showing depletion of quiescent states and emergence of a stress state. (f–o) *Ldlr*^−/−^ and *Apoe*^−/−^ cohorts. (f,g) *Ldlr*^−/−^ contractile subclusters and composition. (h,i) *Apoe*^−/−^ contractile subclusters and composition. (j,k) Primed-marker violins across *Ldlr*^−/−^ (j) and *Apoe*^−/−^ (k) subclusters. (l,m) *Ldlr*^−/−^ module scores and marker dot plot. (n,o) Corresponding *Apoe*^−/−^ analyses.

**Extended Data Fig. 6.**
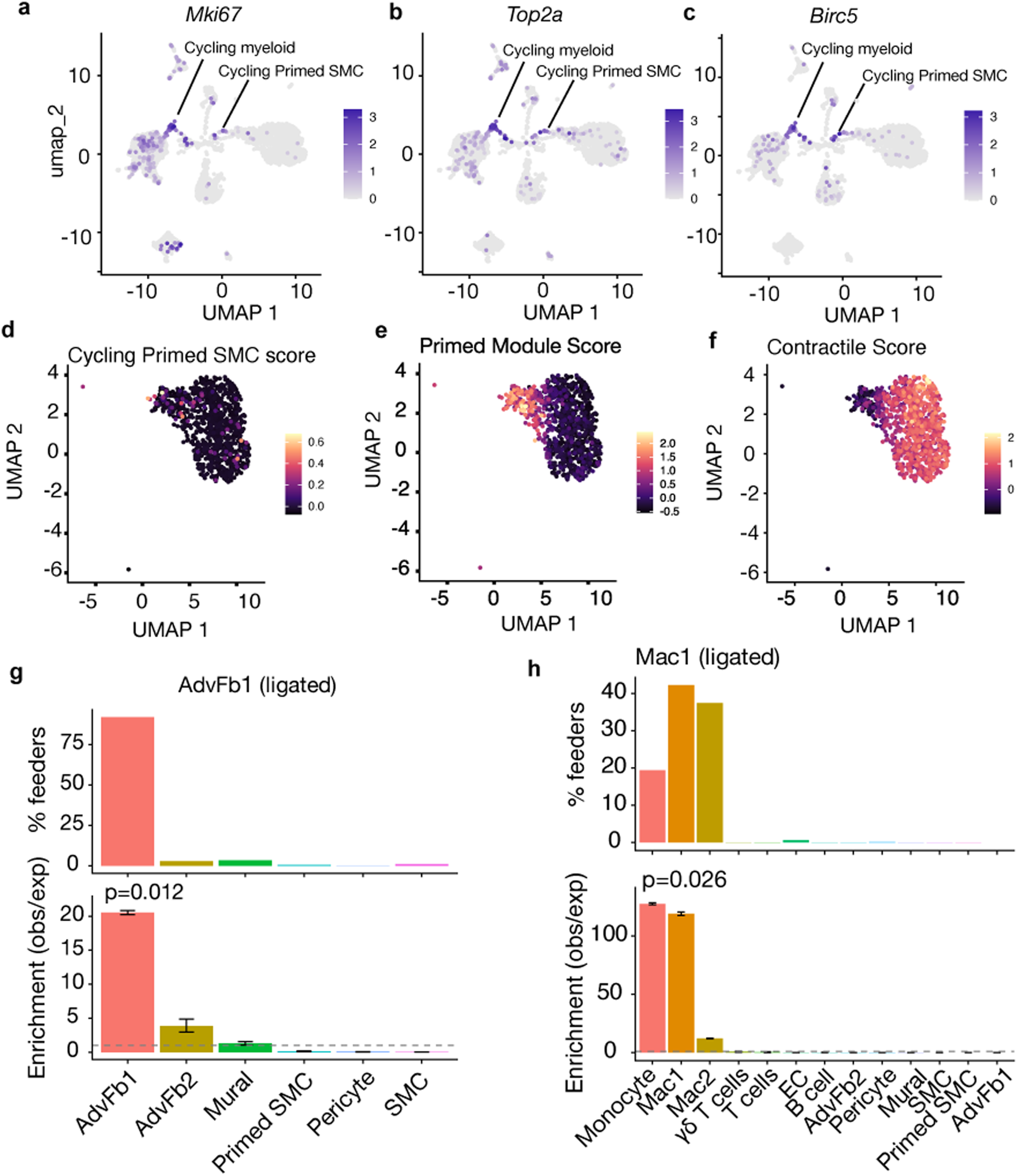
Additional cycling and lineage-specificity controls for carotid ligation. (a–c) Feature plots of Mki67 (a), Top2a (b) and Birc5 (c), resolving Cycling Primed SMC and Cycling myeloid foci. (d–f) Cycling Primed (d), Primed (e) and contractile (f) module scores across the ligated SMC compartment. (g,h) kNN specificity controls using the sham reference: ligated AdvFb1 self-sources from sham AdvFb1 (∼92%; ∼21×; g), while ligated Mac1 maps exclusively to myeloid populations with SMC contributions indistinguishable from zero (h).

**Extended Data Fig. 7.**
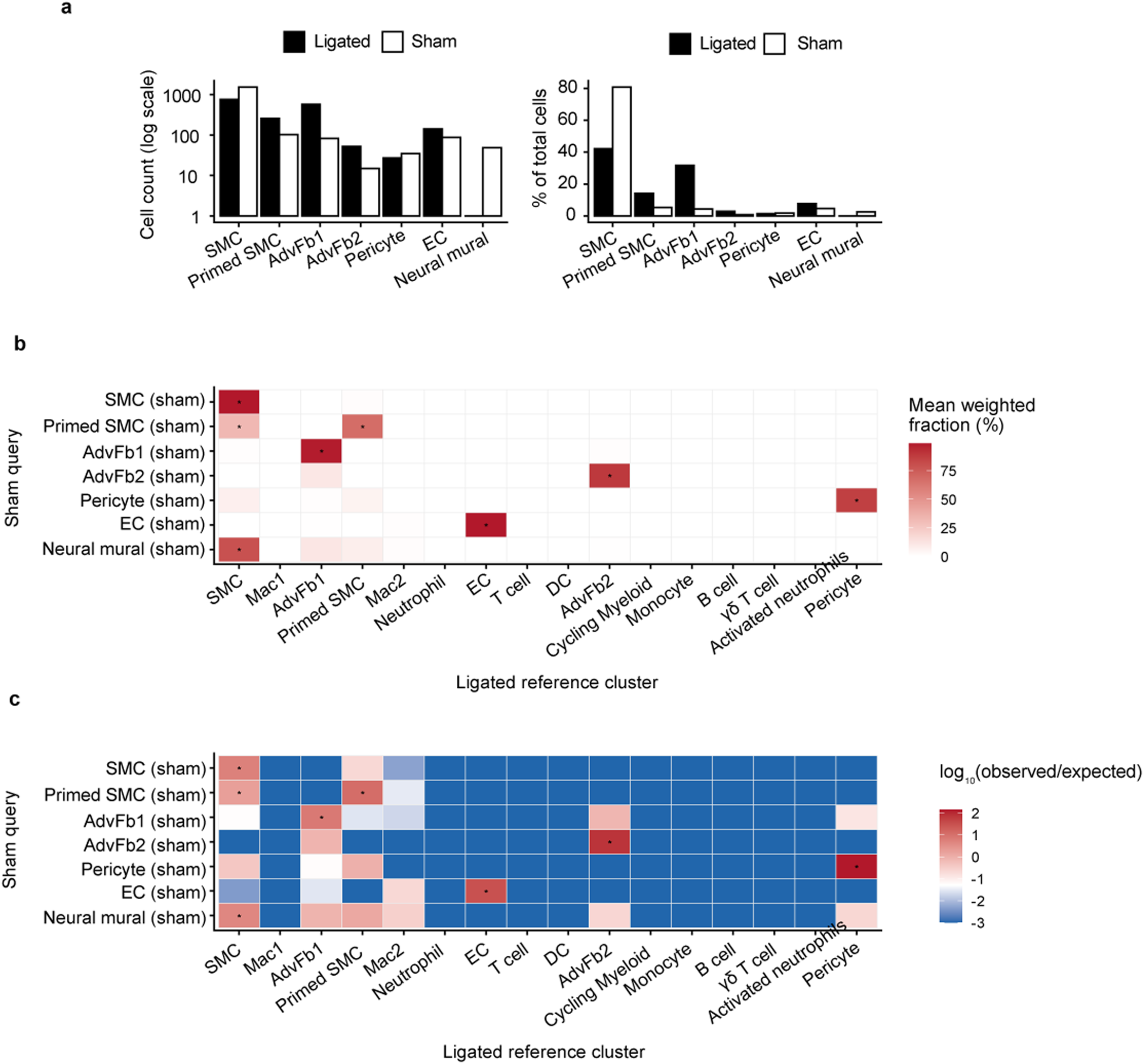
Reciprocal (Sham→Ligated) kNN mapping shows each resident population persists as its own ligated counterpart. (a) Mesenchymal/vascular composition in sham and ligated carotid. (b,c) Reciprocal kNN mapping of each sham population to ligated neighbours: mean weighted neighbour fraction (b) and enrichment over background (c). Each resident population maps principally to its corresponding ligated state. Sham contractile SMCs are depleted as a source of ligated Primed SMCs (0.22×), mirroring Fig. 2; the modest reverse proximity reflects the retained contractile programme rather than directional flux.

**Extended Data Fig. 8.**
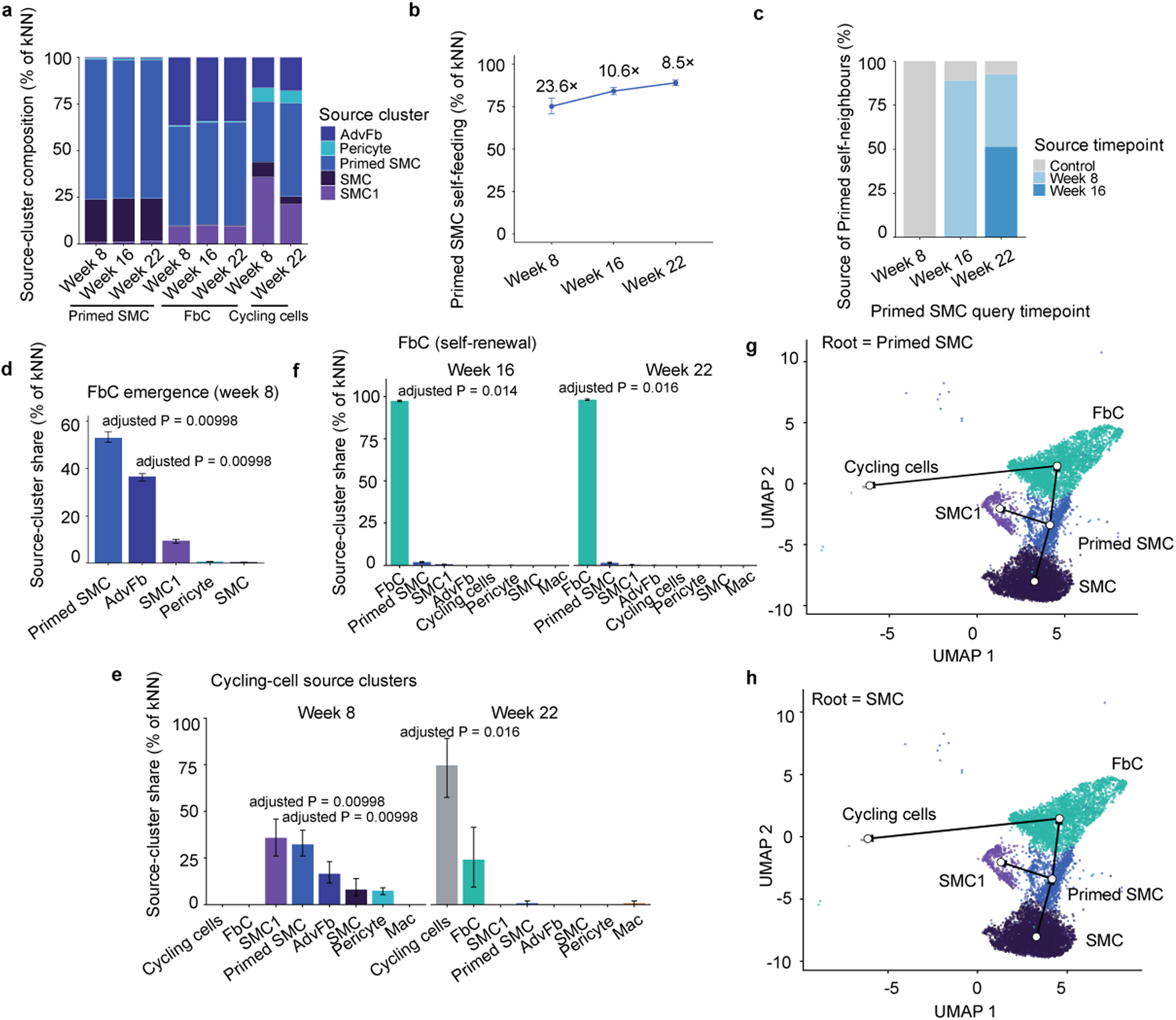
Autonomous expansion and self-renewal of the Primed SMC pool in the ***Apoe*^−/−^** model. Forward kNN feeder-source and dual-rooted Slingshot analysis of lineage-positive (ZsGreen1-WPRE > 3.17) *Apoe*^−/−^ cells (Control, Weeks 8/16/22; k = 15, 20 Harmony dimensions). Bars/points, feeder share; error bars, bootstrap 95% CI (50 iterations); p_adj, 500-permutation test, Bonferroni-corrected. The floor scales with cluster number — 0.00998 (five, Week 8), ≈0.014 (seven, Week 16), ≈0.016 (eight, Week 22) — so the strongest later associations read * (p_adj < 0.05) not **. (a) Feeder composition of each query (Primed SMC, FbC, Cycling cells) by source cluster: Primed queries are self-dominated; FbC draw from Primed SMC and AdvFb; Cycling cells from SMC1, Primed SMC and AdvFb (the SMC1 contribution is a cell-cycle effect; see e). (b) Primed SMC self-feeding >75% at every timepoint (fold-enrichment annotated). (c) Temporal source of Primed self-neighbours: later Primed SMC are seeded by earlier Primed SMC, not contemporaneous recruitment. (d) Emerging FbC (Week 8) fed chiefly by Primed SMC and AdvFb (each p_adj = 0.00998), with minor non-significant SMC1 and negligible SMC/pericyte input. (e) Cycling cells — Week 8: SMC1 and Primed SMC (each p_adj = 0.00998); Week 22: predominantly self-renewing with an FbC contribution (p_adj = 0.016). Cell-cycle-phase regression resolves the Week-8 SMC1 proximity onto AdvFb and Primed SMC and markedly reduces Week-22 self-renewal, identifying both as cell-cycle-embedding effects (Supplementary Fig. 3a,b,d); the Week-22 population is small (n = 10) and k-sensitive. (f) FbC at Weeks 16 and 22 maintained almost entirely by self-neighbours (p_adj = 0.014, 0.016). (g,h) Dual-rooted Slingshot over the *Apoe*^−/−^ SMC-fate compartment (SMC, SMC1, Primed SMC, FbC, Cycling cells; 10 Harmony dimensions): rooting at Primed SMC (g) places both contractile clusters terminal; rooting at SMC (h) forces Primed SMC to an intermediate. Topology only; direction is adjudicated by the feeder analysis (b–f). FbC, fibromyocyte–fibrochondrocyte (C2+C7); SMC1, minor activated-contractile subset; AdvFb, adventitial fibroblast.

**Extended Data Fig. 9.**
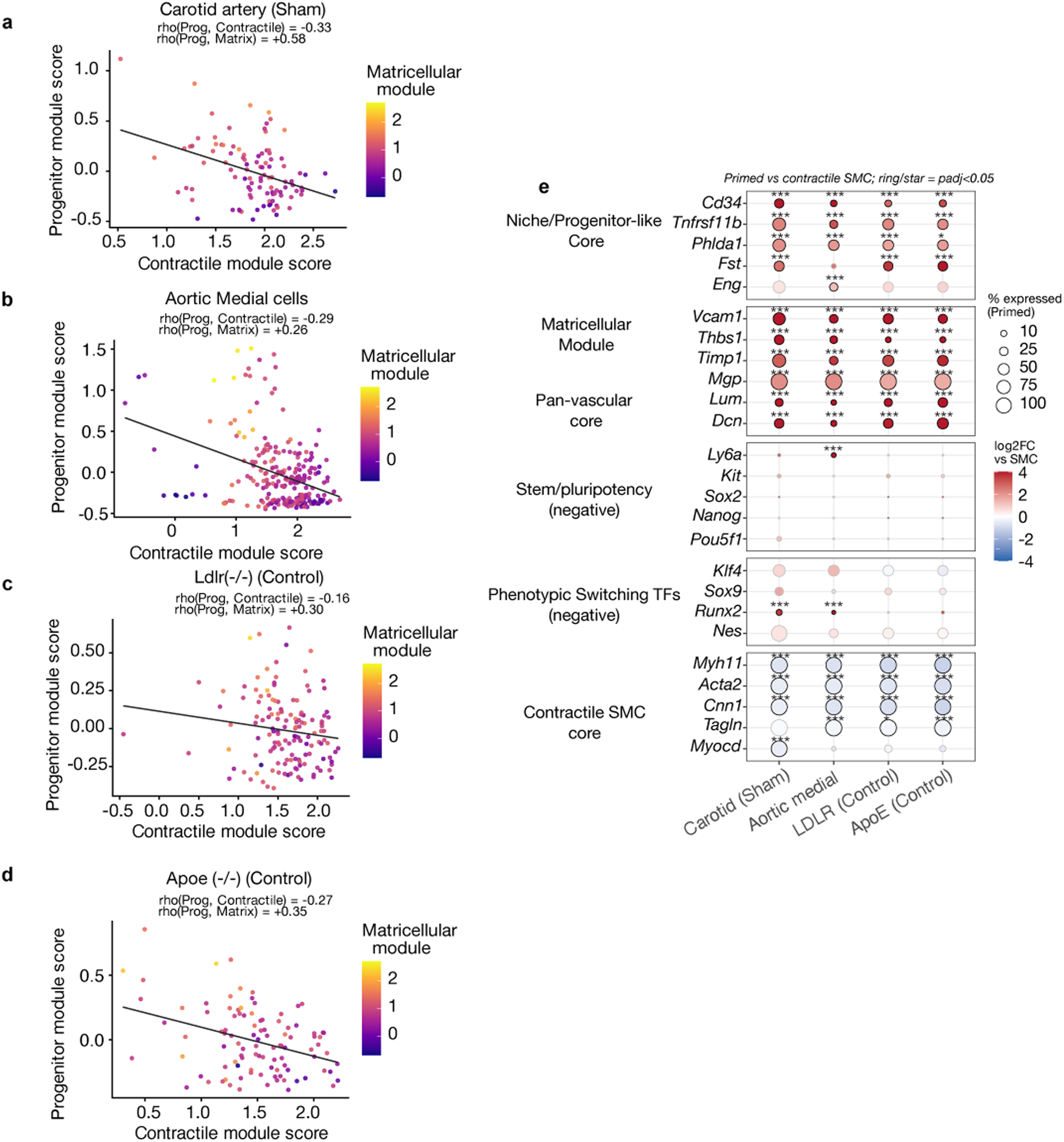
The Primed programme couples progenitor-like and matricellular features on a retained contractile background. (a–d) Per-cell progenitor-versus-contractile module scores coloured by matricellular score in carotid sham (a), healthy aortic media (b), *Ldlr*^−/−^ Control (c) and *Apoe*^−/−^ Control (d). Progenitor and matricellular programmes co-vary and oppose the contractile programme. (e) Primed-versus-contractile log₂ fold changes across progenitor-like, matricellular, pan-vascular, stem/pluripotency, phenotypic-switching and contractile modules. Primed cells enrich the first three programmes and deplete the contractile core; pluripotency and phenotypic-switching markers are not generally enriched.

**Extended Data Fig. 10.**
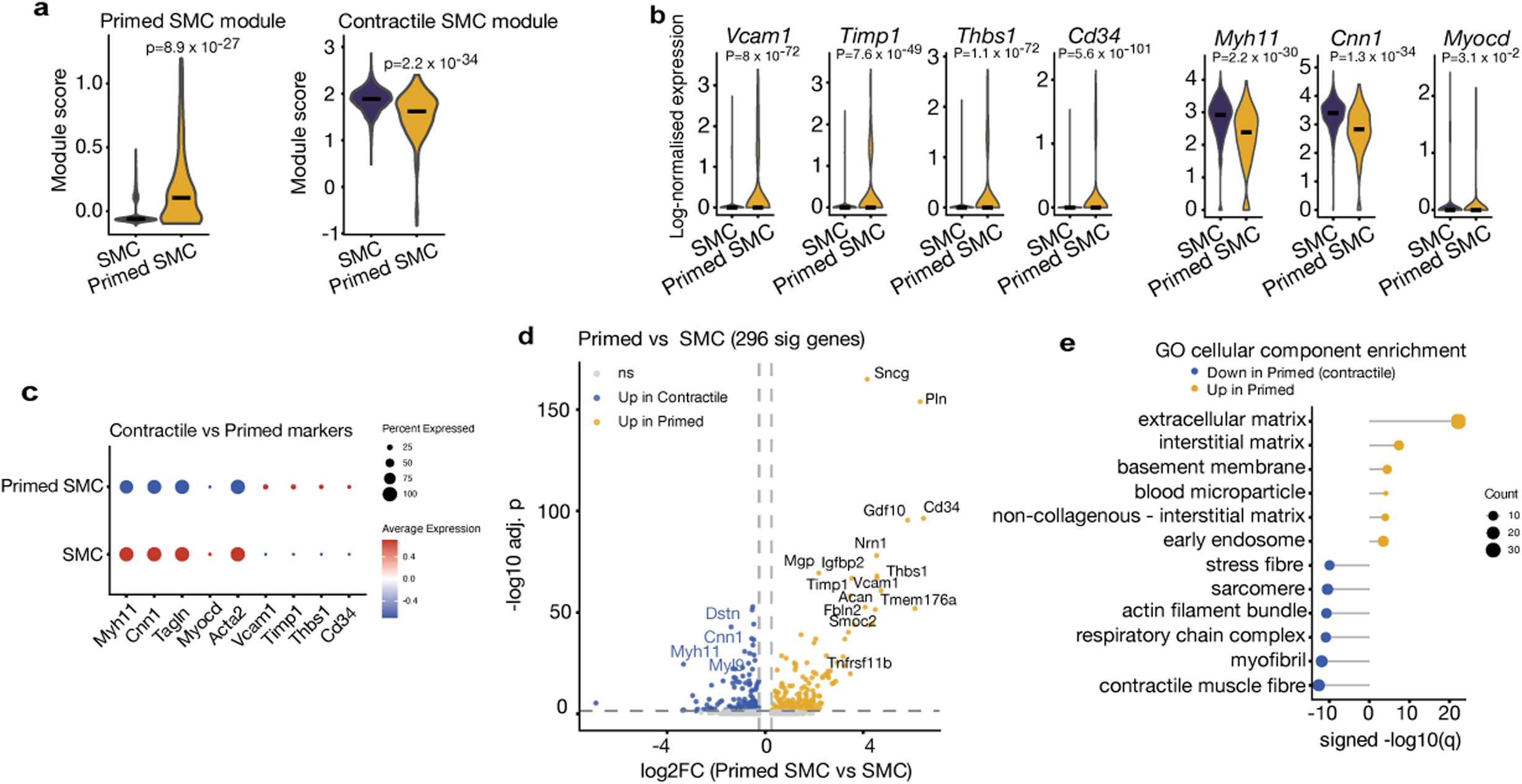
Detailed validation of Primed SMCs in healthy lineage-traced aortic media. FACS-purified Myh11-CreERT² lineage-marked aortic medial cells. (a) Primed and contractile module scores in Primed versus contractile SMC. (b) Violins of Primed and contractile genes. (c) Dot plot of contractile and Primed markers. (d) Volcano plot, Primed versus contractile SMC (296 significant genes; |log₂FC| >0.25, BH-adjusted p <0.05). (e) GO Cellular Component enrichment for genes increased or decreased in Primed SMC.

## Supplementary Figure legends

Supplementary Fig. 1 Ligation-associated reductions in library complexity are concentrated mainly in contractile SMCs.

(a) Whole-atlas QC: UMI count and genes detected, Sham versus Ligated; ligation lowers median UMIs by 45% (P = 1.74 × 10⁻²⁵⁰) and genes by 30% (P = 3.27 × 10⁻²⁶⁴). (b) Per-cluster signed −log₁₀(P) (Ligated versus Sham) for nCount and nFeature: the reduction is driven almost entirely by contractile SMC, with other lineages non-significant or modest (purple, lower in Ligated; amber, higher; ns, P ≥ 0.05; *, P < 0.05; **, P < 0.01; ***, P < 0.001; ****, P < 0.0001). (c) Focused QC: contractile SMC show reduced UMIs (−27%; P = 1.01 × 10⁻¹¹³) and genes (−14%; P = 1.41 × 10⁻¹⁰⁸), whereas Primed SMC are unchanged (UMI +8%, P = 0.179; genes +4%, P = 0.224). Boxes show medians and interquartile ranges; red points show means. P values are raw, unadjusted two-sided cell-level Wilcoxon rank-sum tests. Ten vessels per condition were pooled into one capture, so cells, rather than biological replicates, are the statistical units.

Supplementary Fig. 2 Lineage-tracing recovery, sort-fraction contributions and labelling reproducibility across the ***Ldlr*^−/−^** and ***Apoe*^−/−^**cohorts.

Colour denotes model: Shared (common baseline Control; grey), *Ldlr*^−/−^ (blue) and *Apoe*^−/−^ (orange). (a) Total cells recovered per timepoint and cohort. (b) Cluster-resolved counts (contractile SMC, Primed SMC and FbC) across timepoints, faceted by cohort: contractile SMC decline as Primed SMC and FbC accumulate. (c) Transcript-defined lineage-positive cells (ZsGreen1-WPRE > 3.17) per timepoint and cohort. (d) Contractile SMC, Primed SMC and FbC composition as a fraction of the annotated lineage-positive vascular/mesenchymal cast (SMC, Primed SMC, FbC, pericyte, AdvFb, cycling cells and Mac1); this denominator excludes lineage-positive cells outside that cast. (e) Transcript-defined lineage-positive fraction among all captured cells, stable at approximately 40–48% across cohorts and timepoints. (f) Counts of transcript-defined lineage-positive cells recovered from each FACS-negative (ZsGreen⁻) and FACS-positive (ZsGreen⁺) library by model and time point. FACS-positive fractions yielded substantially more lineage-positive cells, while residual lineage-positive cells in FACS-negative fractions represent gate-missed recombined cells recovered by the transcript threshold.

Supplementary Fig. 3 Cycling cells are the proliferative arm of the Primed→FbC axis; their apparent SMC1 origin and late self-renewal are cell-cycle artefacts, and feeder calls are robust to neighbourhood size.

(a) Control feeder share of Week-8 *Apoe*^−/−^ cycling cells (k = 15, 20 Harmony dimensions; n = 19) before versus after S/G2M-phase regression and re-embedding (500 permutations with Bonferroni correction; **, adjusted P < 0.01): regression abolishes SMC1 (11.1× → 2.1×, not significant) and resolves origin onto AdvFb (→ 7.7×) and Primed SMC (→ 4.0×). (b) Fate of the 32 cycling cells after cell-cycle-corrected re-clustering, by dominant non-cycling identity: predominantly FbC (69%) and Primed SMC (9%). (c,d) Feeder share versus neighbourhood size (k = 5–50) at emergence (solid) and late (dashed) timepoints (dotted line, k = 15). (c) *Ldlr*^−/−^ (Weeks 16 and 26), original embedding: emergence is fed by Primed SMC (approximately 59% for k ≥ 15); the Week-26 self-renewal estimate is strongly k-dependent (approximately 94% → 60% → 19% at k = 5, 15 and 50). (d) *Apoe*^−/−^ (Weeks 8 and 22), cell-cycle-corrected: Week-8 origin is AdvFb plus Primed SMC across all k, with minimal SMC1 contribution; by Week 22, cells map mainly to FbC with only minor self-feeding. Late-timepoint self-renewal rests on small populations (*Ldlr*^−/−^ n = 22; *Apoe*^−/−^ n = 10) and indicates tendencies, not fixed values; the Primed and FbC calls carrying the main conclusions are k-independent.

Supplementary Fig. 4 Integration-free cross-cohort comparisons provide qualified support for the Primed SMC identity.

Cohort fold-change similarity (CFS) scores transcriptome-wide concordance between per-cluster fold-change signatures computed independently within each cohort’s uncorrected RNA assay, without integration or batch correction (values symmetric). Baseline/uninjured cells only (sham carotid; control *Ldlr*^−/−^/*Apoe*^−/−^; healthy aortic medial), lineage-gated where applicable (ZsGreen > 3.17 for *Ldlr*^−/−^/*Apoe*^−/−^; Confetti-sorted medial), restricted to the shared mesenchymal roster. Baseline Primed SMC: carotid n = 101, *Ldlr*^−/−^ n = 141, *Apoe*^−/−^ n = 100, aortic medial n = 240 (SMC identity lineage-confirmed in three cohorts; transcriptionally defined in carotid, 79/101 Myh11 > 3). (a) Cross-cohort Primed↔Primed CFS matrix; boxed = reciprocal-best. Five of six pairs positive, carotid↔aortic media (−1.1) the exception. (b) Each source cohort’s Primed SMC (rows) scored against partner Primed SMC (amber) and partner adventitial fibroblast (green) clusters; FACS-sorted aortic medial source shaded; these lineage-marked SMCs are ECM-dense and can score as fibroblast-like. (c) As (b) under cell-number-balanced downsampling (120 cells per type; robustness control).

**Supplementary Fig. 5 Orthogonal graph-connectivity and temporal-flow analyses identify the Primed SMC as the bridge to the disease-state compartment.** Five mural roles (contractile SMC, Primed SMC, fibromyocyte, fibrochondrocyte, cycling) in the *Ldlr*^−/−^ (DS3) and *Apoe*^−/−^ (DS5) cohorts, on a shared layout. **(a)** Connectivity, no time axis. PAGA-style observed/expected kNN connectivity (abstracted graph, top; symmetric matrix, bottom; shared key 0–0.36). The Primed SMC is the hub linking the contractile pole to the disease states (*Ldlr*^−/−^ Primed–fibrochondrocyte 0.36, Primed–fibromyocyte 0.31; *Apoe*^−/−^ Primed–fibromyocyte 0.24 → fibromyocyte–fibrochondrocyte 0.36). The contractile SMC connects only to the Primed SMC (0.19 / 0.18) and is otherwise inert toward the disease compartment (≤0.08; contractile–fibromyocyte ≈0). (b) Directed flow across time. Temporal-flow enrichment: per adjacent timepoint pair (Control → Wk8 → Wk16 → Wk26 [*Ldlr*^−/−^] / Wk22 [*Apoe*^−/−^]), each later-stage cell’s 30 nearest neighbours among earlier-stage cells were scored by role over 20 dimensions; the matrix gives observed/expected enrichment of each source role (earlier stage; rows) for each target role (later stage; columns), diverging key 1/16–16 (1 = expected). Every disease state is self-sustaining once it emerges (diagonal: cycling 64.6 / 39.1, fibrochondrocyte 12.5 / 9.3, fibromyocyte 4.0 / 2.6), and among the baseline states only the Primed SMC self-renews (5.0 / 5.9); the contractile SMC feeds no disease target (≤0.11 in both cohorts).

Supplementary Fig. 6 Transcript-level lineage definition excludes phagocytic myeloid false-positives across both atherosclerosis cohorts.

SMC lineage is marked by the Myh11-CreERT2–driven ZsGreen reporter, so ZsGreen protein in a non-SMC cell is SMC-lineage–derived by definition. Each cell was cross-classified by FACS sort status and transcript-level reporter detection (ZsGreen1-WPRE > 3.17) into True Lineage (FACS+/transcript+), Discordant (FACS+/transcript−), Gate-missed (FACS−/transcript+) and Negative (FACS−/transcript−). Group sizes (*Ldlr*^−/−^/*Apoe*^−/−^): 11,657/10,699; 3,667/2,599; 299/289; 11,050/10,455. (a–c) *Ldlr*^−/−^; (d–f) *Apoe*^−/−^. (a,d) Discordant cells per Harmony cluster, coloured by provisional identity (SMC, immune, fibroblast, EC; argmax of lineage-marker means): they localise predominantly to immune clusters (*Ldlr*^−/−^ 54.4%, *Apoe*^−/−^ 53.1%) and only marginally to SMC (8.8%, 4.5%). (b,e) Marker dot plot across the four groups (SMC, myeloid/immune, efferocytosis genes and the reporter transcript; dot size, % expressing; colour, scaled mean). True Lineage cells co-express the SMC programme and high reporter transcript, whereas Discordant cells express the myeloid/immune programme with low reporter transcript despite FACS positivity (median ZsGreen1-WPRE — True Lineage 4.38/4.46, Discordant 1.45/1.29, Gate-missed 3.39/3.42, Negative 0.79/0.00). Per cell, 66.0%/69.3% of Discordant cells are myeloid-marker⁺ versus 27.5%/25.1% SMC-marker⁺/immune-marker⁻ (genuine dropout; n = 1,007/653); mean module scores immune +0.70/+0.73, SMC −0.52/−0.58, efferocytosis −0.10/−0.04 — i.e. myeloid cells carrying ingested reporter protein, not an active efferocytosis signature. (c,f) Harmony UMAP with Discordant cells (red) mapping predominantly to myeloid territory

Supplementary Fig. 7 Expression-based definition of the ZsGreen1-WPRE lineage threshold (***Ldlr*^−/−^**reference cohort).

Lineage-positive cells were defined by ZsGreen1-WPRE log-normalised expression > 3.17, set by maximising Youden’s J against the FACS sort gate (n = 26,673; AUC = 0.90, sensitivity = 0.76, specificity = 0.97) and applied globally to both cohorts and all timepoints. (a) Reporter expression in the two FACS fractions (dashed line, > 3.17); the sort-positive fraction contains a substantial sub-threshold population (imperfect protein gating). (b) The same by transcript-defined status at the > 3.17 cutoff. (c) Kernel density across all cells; the distribution is bimodal, with the threshold separating the modes. (d) FACS gate versus transcript threshold: concordant 11,657 (positive) and 11,050 (negative); discordant 3,667 (sort-positive, transcript-negative; predominantly myeloid false-positives with a smaller genuine transcript-dropout component) and 299 (sort-negative, transcript-positive; recovered recombined cells). (e) UMAP of all *Ldlr*^−/−^ cells coloured by reporter expression. (f) The same, lineage-positive cells (> 3.17) coloured by expression; negative cells grey.

