## Supplementary figures and images for "A rare pre-existing progenitor-like Primed SMC compartment is the dominant inferred source of SMC-derived cellularity in vascular injury and atherosclerosis"

### Suppl Fig 1

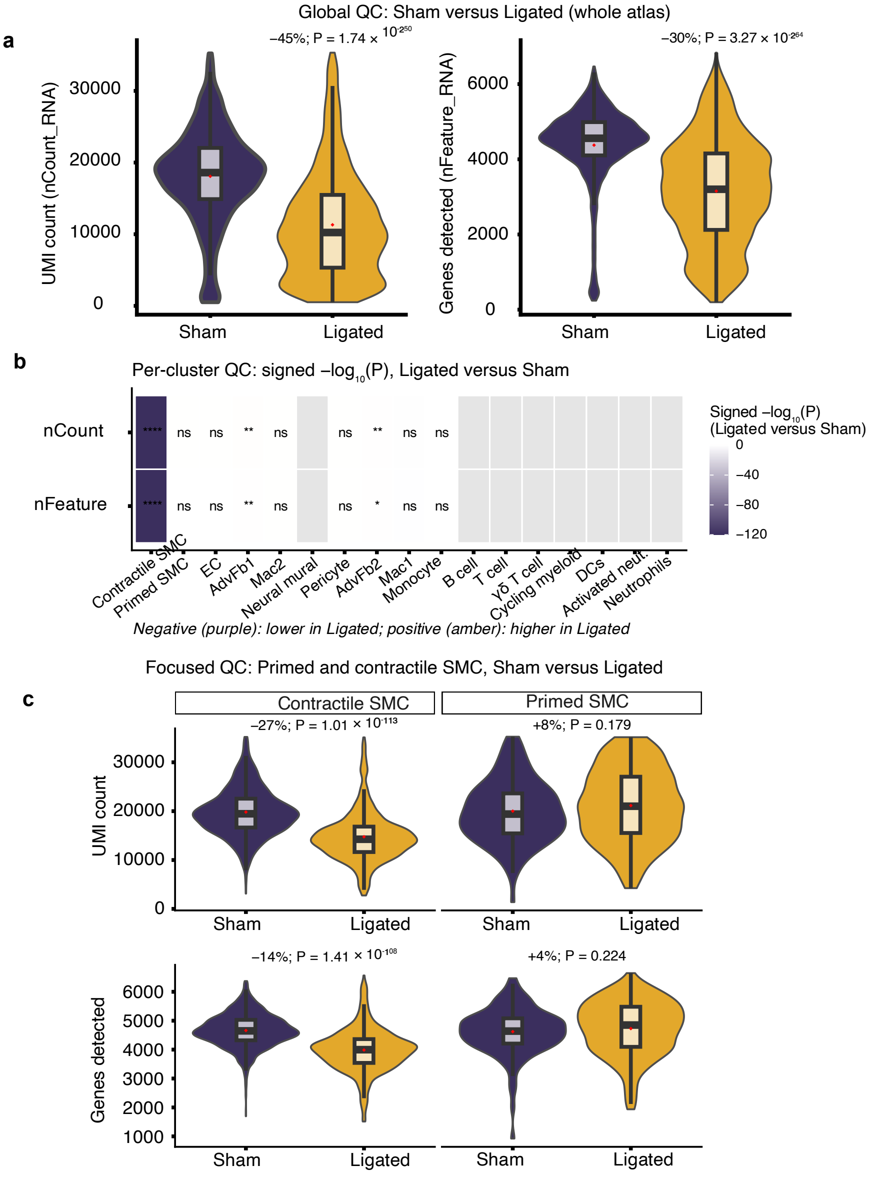

### Suppl Fig 2

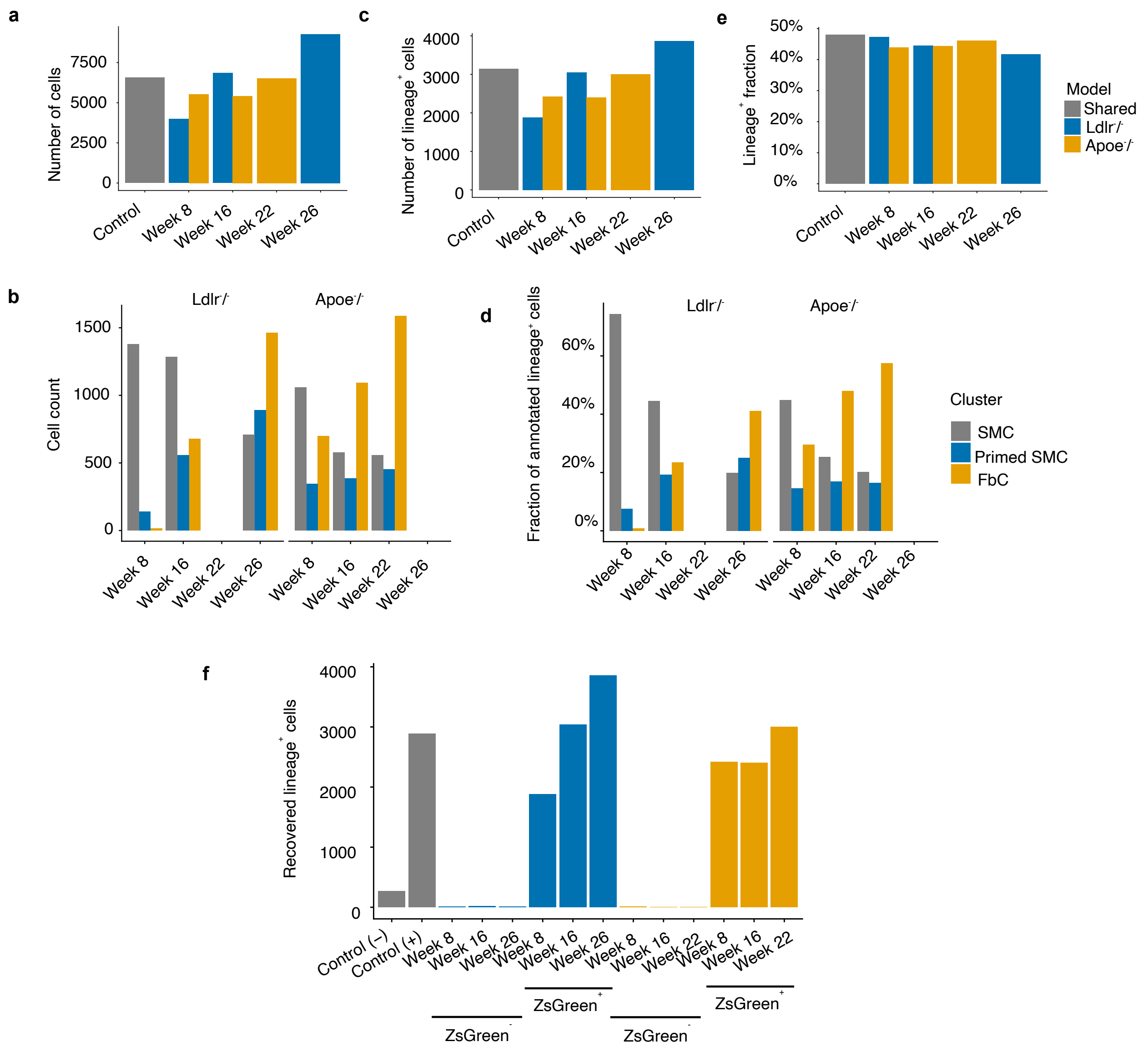

### Suppl Fig 3

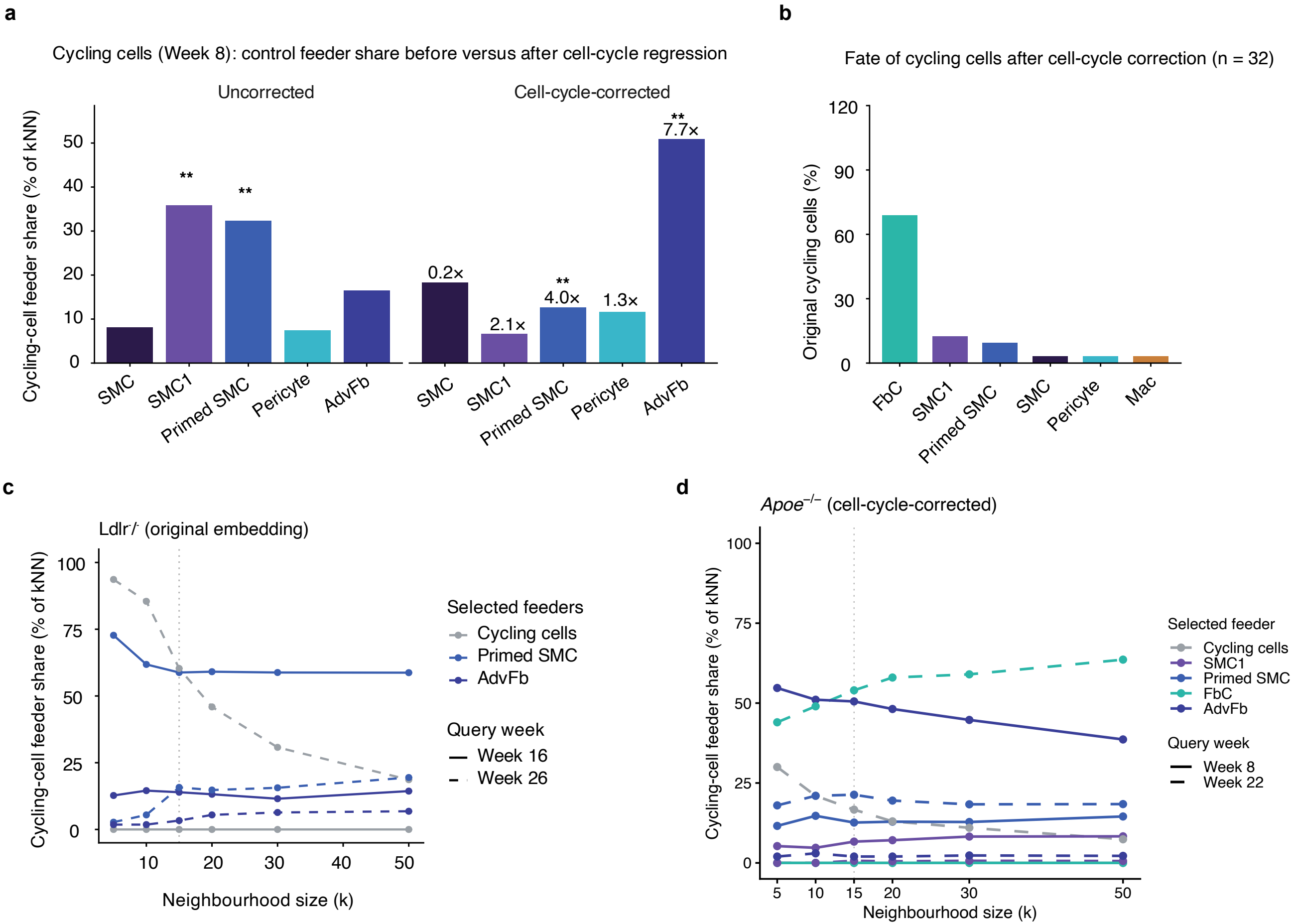

### Suppl Fig 4

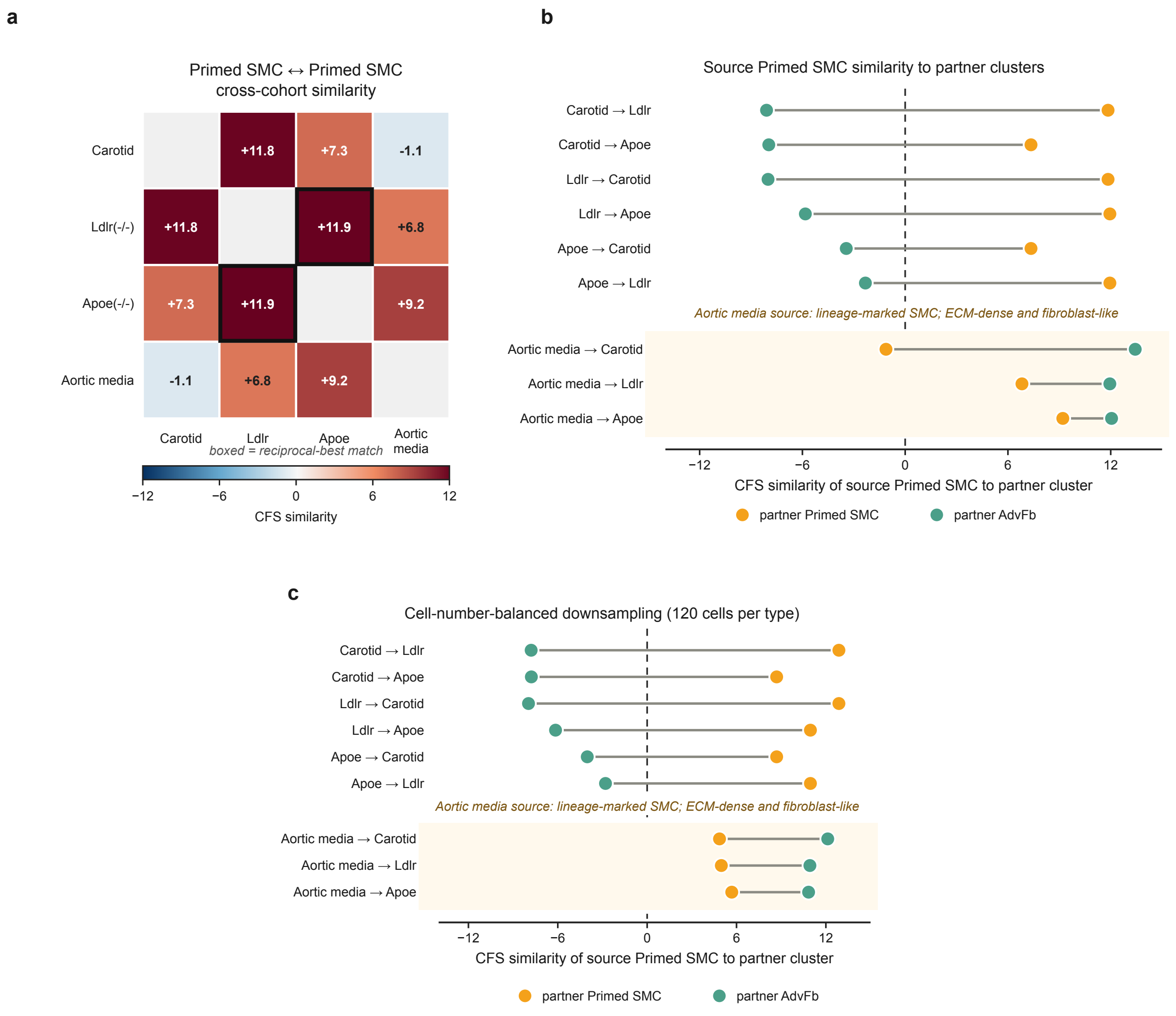

### Suppl Fig 5

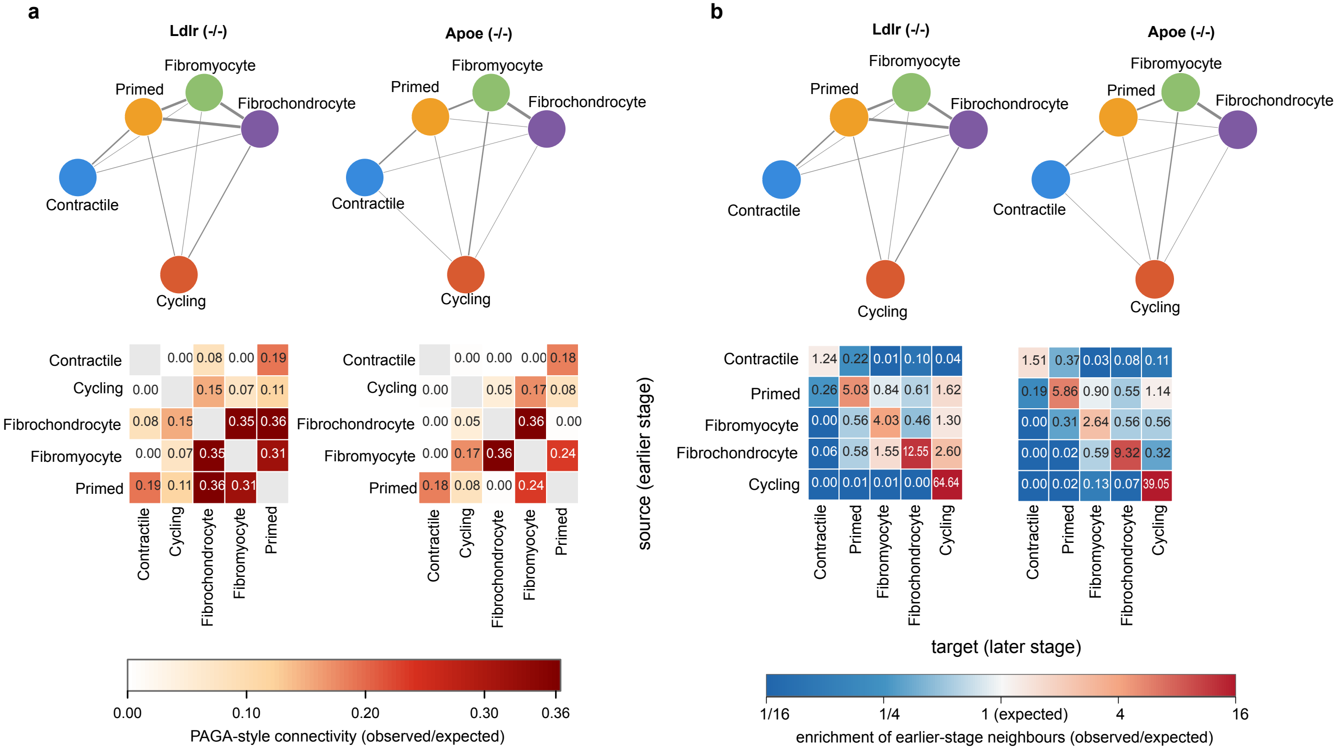

### Suppl Fig 6

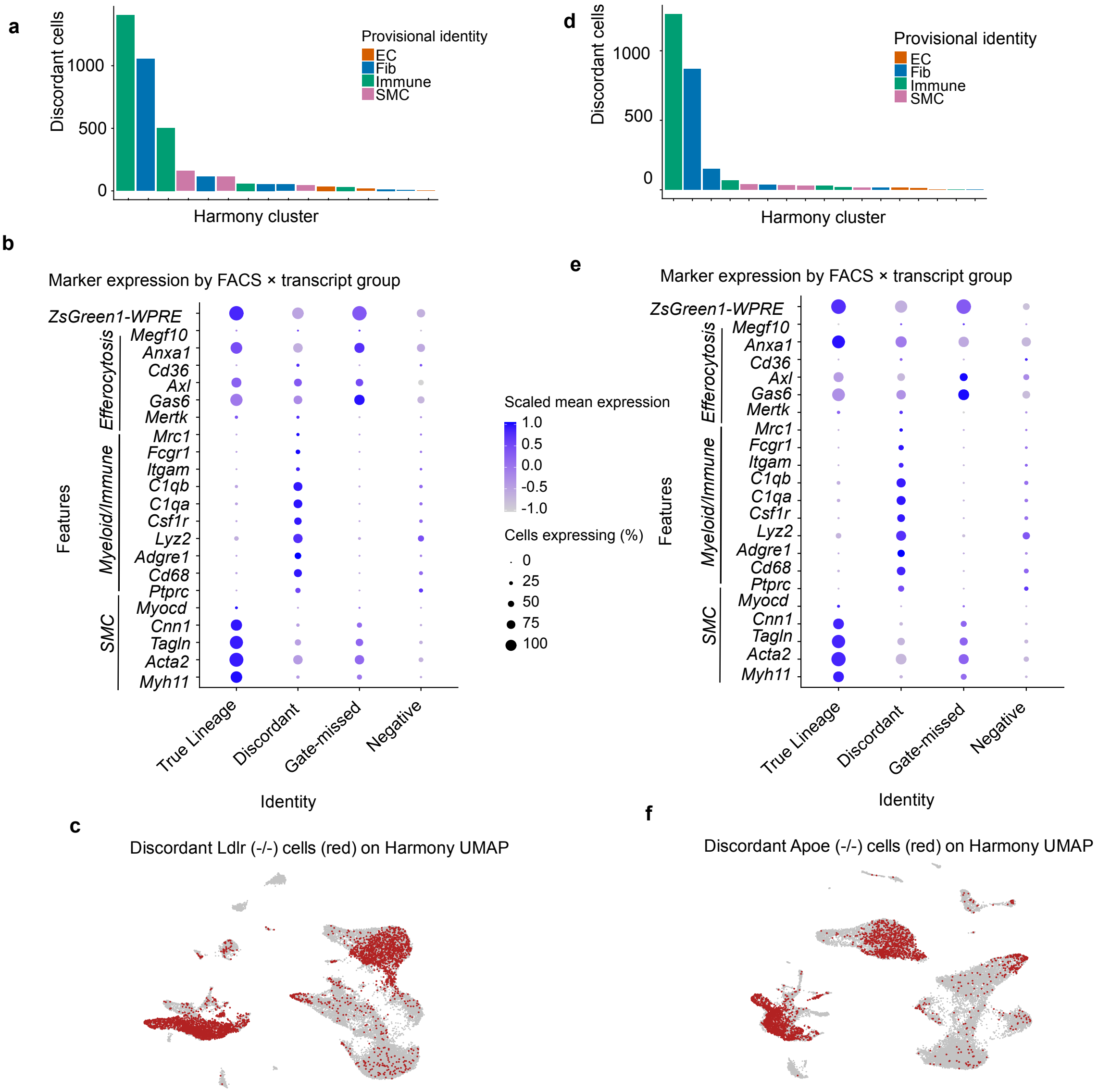

### Suppl Fig 7

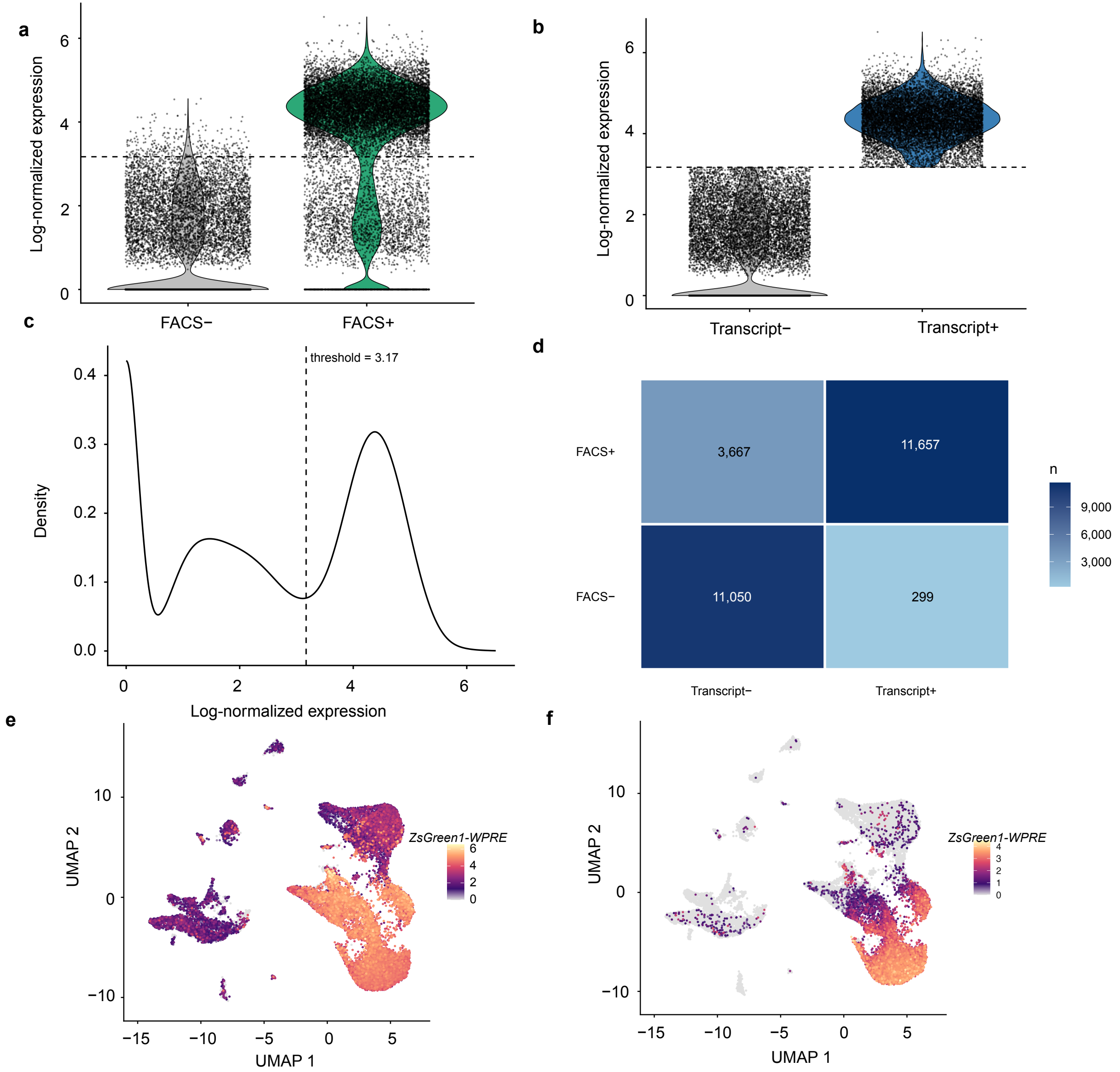
