## Supplementary Table 1 for "A rare pre-existing progenitor-like Primed SMC compartment is the dominant inferred source of SMC-derived cellularity in vascular injury and atherosclerosis"

**Cluster annotation of the carotid artery-ligation single-cell atlas.**

Harmony-integrated clustering of quality-controlled cells from sham and ligated control arms resolved 17 clusters (0–16). Annotations were assigned from canonical marker expression and one-vs-rest differential expression (versus all other clusters). Marker values are mean normalized expression and cell numbers are given per arm; Primed SMC (C3) is shown for two contrasts: versus all other clusters, and versus contractile SMC (C0).

| **Cluster** | **Annotation** | **Key markers (mean normalized expression)** | **Sham** | **Ligated** |
| --- | --- | --- | --- | --- |
| 0 | SMC | *Acta2* 236, *Tagln* 142.5, *Myh11* 44.4, *Cnn1* 12.8, *Myocd* 4.1 | 1,534 | 761 |
| 1 | Mac1 | *Apoe* 60.7, *Lyz2* 75.9, *C1qb* 19.6, *C1qa* 17.8, *Mrc1* 4.4, *Trem2* 2.6 | 7 | 1,229 |
| 2 | AdvFb1 | *Dcn* 90.6, *Lum* 40.9, *Serpinf1* 19, *Clec3b* 6.7, *Pi16* 7.3, *Pdgfra* 4.1 | 81 | 572 |
| 3 | Primed SMC *(vs all clusters)* | *Mgp* 318.3, *Spp1* 80.1, *Eln* 72.2, *Lum* 12.9, *Tnc* 1.3, *Nrn1* 0.8, *Acan* 0.7 | 101 | 255 |
|  | Primed SMC *(vs contractile SMC, C0)* | *Timp1* 7.9, *Thbs1* 6.4, *Cxcl12* 5.1, *Vcam1* 4.5, *Ccnd1* 0.6, *Cd34* 0.5 (Myh11 retained, 16.1 vs 44.4 in C0) |  |  |
| 4 | Mac2 | *Apoe* 28.8, *Lyz2* 27.3, *C1qb* 8.9 | 61 | 180 |
| 5 | Neutrophil | *S100a9* 248.9, *S100a8* 169.1, *Csf3r* 16.9, *Mmp9* 13.3, *Cxcr2* 7 | 0 | 410 |
| 6 | Endothelium | *Pecam1* 19, *Cdh5* 5.4, *Cldn5* 5.2, *Tie1* 2.3, *Sox18* 2.2, *Sox17* 1.9 | 87 | 140 |
| 7 | T cell | *Cd3d* 6.3, *Cd3e* 3.2, *Trac* 3.2, *Lef1* 4.3, *Cd2* 2.7 | 1 | 221 |
| 8 | Dendritic cell | *Ccr7* 3.5, *Cd209a* 2.8, *Clec10a* 2.1, *Flt3* 1.6 | 0 | 153 |
| 9 | AdvFb2 | *Dcn* 69.1, *Lum* 24, *Timp1* 6.1, *Serpinf1* 16.1, *Pi16* 5.7, *Ly6a* 5.5 | 14 | 51 |
| 10 | Cycling (myeloid) | *Top2a* 7.4, *Mki67* 6.2, *Birc5* 4.1, *Ccna2* 1.6 | 0 | 70 |
| 11 | Monocyte | *Csf1r* 5.9, *Itgal* 7, *Adgre4* 4, *Fcgr4* 3.2, *Cd300e* 1.6 | 3 | 89 |
| 12 | B cell | *Cd79a* 22, *Cd79b* 8.7, *Ms4a1* 7.4, *Pax5* 3.9, *Cd19* 1.2 | 1 | 64 |
| 13 | Neural Mural (neural-crest, Dlx+) | *Acta2* 72.9, *Tagln* 47.5, *Myh11* 30.2, *Dlx5* 1.1, *Dlx6* 1.5, *Shox2* 1.1 | 48 | 0 |
| 14 | γδ T cell | *Trdc* 14.5, *Il17a* 7.3, *Cd3d* 7, *Il23r* 6.1, *Trdv4* 3 | 1 | 41 |
| 15 | Activated neutrophil (ISG+) | *Rsad2* 53.2, *Cxcl10* 40, *Ifit1* 28.7, *Ifit3* 28.9, *S100a9* 418.6 | 0 | 10 |
| 16 | Pericyte | *Rgs5* 26.3, *Notch3* 8.6, *Ndufa4l2* 3.1, *Pdgfrb* 2.9, *Higd1b* 0.6 | 34 | 26 |
