## Supplementary Table 2 for "A rare pre-existing progenitor-like Primed SMC compartment is the dominant inferred source of SMC-derived cellularity in vascular injury and atherosclerosis"

**Cluster annotation of the Ldlr⁻/⁻ atherosclerosis Myh11-lineage single-cell atlas.**

Harmony-integrated clustering of Myh11-CreERT²; ZsGreen1-WPRE lineage-traced aortic cells (Pan et al., Ldlr⁻/⁻ cohort) across Control, Western-diet Week 8, Week 16 and Week 26. Lineage-positive cells were defined as ZsGreen1-WPRE > 3.17 (log-normalised expression), identifying 11,956 lineage-positive cells (concordant with the dataset’s native lineage annotation to within one cell). Annotations were assigned from canonical markers and one-vs-rest differential expression among lineage-positive clusters. Cell numbers are lineage-positive cells per timepoint; “Lineage⁺ (%)” is the ZsGreen-positive fraction of the parent Harmony cluster(s). Primed SMC is shown for two contrasts: versus all other lineage-positive clusters, and versus contractile SMC in Control. Low-quality and predominantly lineage-negative clusters (endothelial, dendritic/monocytic, Schwann and unassigned populations) were resolved but excluded from the lineage-positive analysis.

| **Cluster** | **Annotation** | **Lineage⁺ (%)** | **Key markers** | **Control** | **Week 8** | **Week 16** | **Week 26** | **Total** |
| --- | --- | --- | --- | --- | --- | --- | --- | --- |
| 0 | SMC | 94.1 | *Myh11*, *Acta2*, *Tagln*, *Cnn1*, *Myocd* | 2,435 | 1,378 | 1,286 | 710 | **5,809** |
| 4 | Primed SMC (vs all clusters) | 95.1 | *Serpina3n*, *Cp*, *Sncg*, *Vcam1*, *Cxcl12*, *Tm4sf1*, *Lrg1* | 141 | 140 | 556 | 890 | **1,727** |
|  | Primed SMC (vs contractile SMC, Control) |  | *Spp1*, *Vcam1*, *Timp1*, *Cxcl12*, *Lum*, *Dcn* (Primed module Vcam1/Timp1/Thbs1/Cd34; Cd34 +2.86 log₂FC, p_adj 6.0×10⁻⁸) |  |  |  |  |  |
| 2 | FbC (fibrochondrocyte) | 79.3 | *Cytl1*, *Spp1*, *Thbs1*, *Crabp2*, *Ecrg4*, *Cdkn2a* | 3 | 15 | 679 | 1,464 | **2,161** |
| 5 | Pericyte/Neural Mural | 68.5 | *Rgs5*, *Notch3*, *Dlx5*, *Dlx6*, *Tfap2b*, *Msx2* | 288 | 298 | 293 | 319 | **1,198** |
| 1, 6, 16 | AdvFb | 6.4 | *Dcn*, *Lum*, *Serpinf1*, *Dpt*, *Clec3b*, *Pdgfra* | 196 | 22 | 53 | 140 | **411** |
| 18 | Cycling cells | 43.4 | *Mki67*, *Top2a*, *Ccnb2*, *Birc5*, *Cdca3* | 0 | 0 | 11 | 22 | **33** |
| 3 | Mac (lineage⁺ macrophage-like) | 1.0 | *C1qa*, *C1qb*, *Lyz2*, *Tyrobp*, *Ctss* | 0 | 0 | 9 | 14 | **23** |
|  | **Total lineage⁺ (analysed cells)** |  |  | **3,063** | **1,853** | **2,887** | **3,559** | **11,362** |

*Marker genes are leading identity markers from the one-vs-rest test; for Primed SMC, the second row lists genes enriched versus contractile SMC in Control (the Primed module plus matricellular genes). Cd34 marks a subset of Primed SMC (versus contractile, Control: avg log₂FC +2.86, p_adj 6.0×10⁻⁸); Thbs1 lies below the 10% detection threshold in Control and is captured by the dot plot rather than the differential test while remaining a defining module gene (module score p_adj 1.04×10⁻³³; Fig. 3H). AdvFb (clusters 1/6/16) and Mac (cluster 3) are predominantly lineage-negative populations; only their minor ZsGreen-positive subsets are tabulated. The Pericyte cluster (5) also expresses neural-crest mural markers (Dlx5, Dlx6, Tfap2b, Msx2).*
