## Supplementary Table 3 for "A rare pre-existing progenitor-like Primed SMC compartment is the dominant inferred source of SMC-derived cellularity in vascular injury and atherosclerosis"

**Apoe^−/−^ lineage-positive cluster composition across disease progression.**

| **Cell type** | **Cluster(s)** | **Key markers** | **Control** | **Week 8** | **Week 16** | **Week 22** | **Total (% of lineage^+^)** |
| --- | --- | --- | --- | --- | --- | --- | --- |
| Contractile SMC | 0 | Myh11, Cnn1, Tagln, Acta2, Myocd | 2,459 (78.2%) | 1,059 (43.7%) | 579 (24.1%) | 557 (18.6%) | 4,654 (42.4%) |
| SMC1 (activated contractile) | 10 | Acta2, Tagln, Myh11; Lcn2, Saa3, Ly6a, Spp1, Mmp3 | 101 (3.2%) | 69 (2.8%) | 123 (5.1%) | 235 (7.8%) | 528 (4.8%) |
| Primed SMC | 4 | Vcam1, Timp1, Thbs1, Cd34, Lum, Dcn | 100 (3.2%) | 344 (14.2%) | 386 (16.1%) | 453 (15.1%) | 1,283 (11.7%) |
| FbC (fibromyocyte–fibrochondrocyte) | 2, 7 | Col1a1, Lum, Dcn; Acan, Sox9, Comp, Col2a1 | 0 (0%) | 697 (28.7%) | 1,094 (45.6%) | 1,587 (53.0%) | 3,378 (30.8%) |
| Pericyte | 5 | Notch3, Rgs5, Kcnj8, Pdgfrb | 276 (8.8%) | 220 (9.1%) | 184 (7.7%) | 98 (3.3%) | 778 (7.1%) |
| AdvFb (adventitial fibroblast) | 1, 6 | Pdgfra, Dcn, Lum, Gsn | 209 (6.6%) | 17 (0.7%) | 29 (1.2%) | 29 (1.0%) | 284 (2.6%) |
| Cycling cells | 15 | Mki67, Top2a, Birc5 | 0 (0%) | 19 (0.8%) | 3 (0.1%) | 10 (0.3%) | 32 (0.3%) |
| Macrophage | 3 | Cd68, Lyz2, C1qa, Adgre1 | 0 (0%) | 0 (0%) | 1 (0.04%) | 28 (0.9%) | 29 (0.3%) |
| **Total lineage⁺** |  |  | **3,145** | **2,425** | **2,399** | **2,997** | **10,966** |

FbC = merged clusters 2 + 7; AdvFb = merged clusters 1 + 6. Counts are cell numbers within each timepoint; percentages in parentheses are within-timepoint composition, and the final column gives each population’s share of all lineage-positive cells. Lineage-positive cells were defined by ZsGreen1-WPRE > 3.17 log-normalised counts. Clusters excluded as non-SMC-lineage (lineage-positive contaminants or doublets): endothelium (8, 14), dendritic cell (9), monocyte (11), granulocyte (12), T cell (13), Schwann (16) and mesothelial (17).
