## Supplementary Table 4 for "A rare pre-existing progenitor-like Primed SMC compartment is the dominant inferred source of SMC-derived cellularity in vascular injury and atherosclerosis"

**Module gene-set definitions.**

| **State** | **Module** | **Gene set** |
| --- | --- | --- |
| **Primed SMC** | Primed core — module score (Figs 1E,J, 2D, 3H, 5G, 6F, 7A; Extended Data Figs 3b, 5b, 10a) | *Vcam1, Timp1, Thbs1, Cd34* |
| **Primed SMC** | Primed programme marker panel (Extended Data Fig. 1a) | Progenitor/niche: Cd34, Cxcl12, Fst, Tnfrsf11b; Developmental: Nrn1, Gdf10, Scx; Proliferative: Ccnd1 |
| **Primed SMC** | Primed — sub-cluster marker panel (Extended Data Fig. 5c) | Vcam1, Timp1, Thbs1, Cd34, Ccnd1 |
| **Contractile SMC** | Strict contractile — module score (Figs 1E, 2D; Extended Data Figs 5b, 10a) | *Myh11, Cnn1, Myocd, Carmn* |
| **Contractile SMC** | Extended contractile — sub-cluster marker panel (Extended Data Fig. 5c) | *Myh11, Cnn1, Myocd, Carmn, Tagln, Acta2, Lmod1* |
| **Cycling** | Cycling — module score (Fig. 2D; Extended Data Fig. 6d) | *Mki67, Top2a, Birc5* |
| **FbC (fibrochondrocyte)** | Fibrochondrocyte — module score (Fig. 7B) | *Acan, Sox9, Col2a1, Comp, Chad, Ibsp* |
| **Bridge (Primed → FbC)** | Bridge — marker module (Fig. 7D) | *Spp1, Prg4* |
| **Myeloid** | Immune gating — not displayed | *Cd68, Adgre1, Lyz2, Csf1r, C1qa (doublet removal: Ptprc, Lyz2, C1qa)* |
| **Contractile SMC (stress sub-state)** | Stress — module score and sub-cluster marker panel (Extended Data Fig. 5b,c) | *Atf3, Egr1, Jun, Fos, Fosb, Hspa1a, Hspa1b, Hsph1, Ddit3, Jund* |
| **Primed SMC** | Niche/progenitor-like core — module score and marker panel (Extended Data Fig. 9a–e) | *Cd34, Tnfrsf11b, Phlda1, Fst, Eng* |
| **Primed SMC** | Matricellular — module score and marker panel (Extended Data Fig. 9a–e) | *Vcam1, Thbs1, Timp1* |
| **Pan-vascular** | Pan-vascular core — marker panel (Extended Data Fig. 9e) | *Mgp, Lum, Dcn* |
| **Negative control** | Stem/pluripotency — marker panel (Extended Data Fig. 9e) | *Ly6a, Kit, Sox2, Nanog, Pou5f1* |
| **Negative control** | Phenotypic-switching transcription factors — marker panel (Extended Data Fig. 9e) | *Klf4, Sox9, Runx2, Nes* |
| **Contractile SMC** | Contractile SMC core — module score and marker panel (Extended Data Fig. 9a–e) | *Myh11, Acta2, Cnn1, Tagln, Myocd* |
| **Human SMC** | Primed — module score (Fig. 8C,D,H) | *VCAM1, TIMP1, THBS1, CD34, ENG* |
| **Human SMC** | Contractile — module score for the Primed–contractile axis (Fig. 8C) | *MYH11, MYOCD, CNN1, CARMN, TAGLN, ACTA2, LMOD1, MYL9* |
| **Human SMC** | Contractile identity — module score (Fig. 8I) | *MYH11, CNN1, MYOCD, CARMN* |
| **Human SMC** | Primed–contractile axis (Fig. 8C) | *Per-cell Primed module score minus eight-gene contractile module score* |
| **Human SMC** | Cross-species concordance set (Fig. 8G) | *VCAM1, TIMP1, THBS1, CD34, ENG, TNFRSF11B, FST, PHLDA1, MGP, LUM, DCN* |

**Notes**

Module scores were calculated with Seurat AddModuleScore and are background-corrected relative values (Methods). Cycling Primed SMCs were defined by S or G2/M phase assignment together with detectable expression (counts > 0) of each of Mki67, Top2a and Birc5, not by the cycling module score. Myeloid gating genes were used for the negative-control Mac1 feeder analysis and for aortic-media doublet removal and are not displayed in any figure panel.
